# High-throughput functional characterization of visceral afferents by optical recordings from thoracolumbar and lumbosacral dorsal root ganglions

**DOI:** 10.1101/2021.01.20.427516

**Authors:** Zichao Bian, Tiantian Guo, Shaowei Jiang, Longtu Chen, Jia Liu, Guoan Zheng, Bin Feng

## Abstract

Functional understanding of visceral afferents is important for developing new treatment to visceral hypersensitivity and pain. The sparse distribution of visceral afferents in dorsal root ganglions (DRGs) has challenged conventional electrophysiological recordings. Alternatively, Ca^2+^ indicators like GCaMP6f allow functional characterization by optical recordings. Here we report a turnkey microscopy system that enables simultaneous Ca^2+^ imaging at two parallel focal planes from intact DRG. By using consumer-grade optical components, the microscopy system is cost effective and can be made broadly available without loss of capacity. It records low-intensity fluorescent signals at a wide field of view (1.9 x 1.3 mm) to cover a whole mouse DRG, with a high pixel resolution of 0.7 micron/pixel, a fast frame rate of 50 frames/sec, and the capability of remote focusing without perturbing the sample. The wide scanning range (100 mm) of the motorized sample stage allows convenient recordings of multiple DRGs in thoracic, lumbar, and sacral vertebrae. As a demonstration, we characterized mechanical neural encoding of visceral afferents innervating distal colon and rectum (colorectum) in GCaMP6f mice driven by VGLUT2 promotor. A post-processing routine is developed for conducting unsupervised detection of visceral afferent responses from GCaMP6f recordings, which also compensates the motion artefacts caused by mechanical stimulation of the colorectum. The reported system offers a cost-effective solution for high-throughput recordings of visceral afferent activities from a large volume of DRG tissues. We anticipate a wide application of this microscopy system to expedite our functional understanding of visceral innervations in both health and diseases.

## Introduction

Sensory information from the internal visceral organs is conveyed by visceral afferents, which transduce stimuli into trains of action potentials at the distal nerve endings embedded in visceral tissues. Evoked action potentials are then transmitted to the spinal cord via long axons distal and central to their somata in the dorsal root ganglion (DRG). In pathophysiological conditions, visceral afferents can undergo functional changes to drive the persistence of disease conditions (Anand et al., 2007). For example, the sensitization of afferents innervating distal colon and rectum (colorectum) appears necessary for the prolonged visceral hypersensitivity and pain in irritable bowel syndrome (Feng et al., 2012b). A better functional understanding of visceral afferents in both health and diseases can potentially lead to the development of new treatment methods to reverse visceral hypersensitivity, the management of which is an unmet clinical need (Chen et al., 2017). Functional recordings from visceral afferents are challenged by the sparse nature of visceral innervations, i.e., visceral afferent somata being the minority in the DRG. For example, colorectal afferents make up less than 10% of the total afferent neurons in mouse L6 DRG, and the proportion is much smaller in adjacent DRGs (Guo et al., 2019).

The sparse distribution of visceral afferents in the DRG has prevented a wider application of conventional electrophysiological recordings to characterize visceral afferent functions. Only a handful of reports implemented intracellular DRG recordings by liquid-filled glass electrodes to characterize the neural encoding of afferents innervating the colon (Malin et al., 2011;Hibberd et al., 2016) and stomach (Bielefeldt et al., 2006). This is in contrast to a larger number of studies using similar approaches to record afferents innervating the skin, e.g., (Woodbury and Koerber, 2007;Jankowski et al., 2010;Koerber et al., 2010;Molliver et al., 2011; Jankowski et al., 2012;Vrontou et al., 2013). Alternatively, recordings of visceral afferents were conducted by manually splitting afferent nerve trunk into microns thick filaments for single-fiber recordings (e.g., (Feng and Gebhart, 2015)) or using a miniature suction electrode to record from the nerve surface (e.g., (Peiris et al., 2011)). However, visceral organs are predominantly innervated by unmyelinated C-fibers and thinly myelinated Aδ-fibers (Sengupta and Gebhart, 1994;Danuser et al., 1997;Feng et al., 2012b;Herweijer et al., 2014;Schwartz et al., 2016), and their small axonal diameter has challenged single-fiber recordings using conventional electrodes or electrode arrays. In fact, there has been no convincing evidence in the literature demonstrating successful single-fiber recordings from unmyelinated C-type peripheral axons in mammalians by commercially available electrode arrays. Overall, electrophysiological approaches to characterize visceral afferent functions are technically challenging and can usually report no more than 100 neurons per study in the literature.

Alternatively, the fluorescent Ca^2+^ indicators like the Furo-2 and Flura-4 have allowed the measurement of intracellular calcium concentrations, and corresponding algorithms have been developed to infer neural spike trains from intracellular calcium imaging data (Pachitariu et al., 2018). In addition, genetically encoded calcium indicators (GECI) can be selectively expressed in target neural populations to allow focused functional studies. Recently developed GECIs like GCaMP6f can rapidly alter their fluorescent responses within milliseconds to changes in intracellular calcium concentrations, which has made possible to resolve individual spikes from GCaMP6f calcium responses when using fast scanning imaging methods like the confocal and two-photon microscopy (Podor et al., 2015). Recently, we and others have shown that conventional epi-fluorescence imaging is also capable to resolve individual spikes in CGaMP6f recordings (Emery et al., 2016;Kim et al., 2016;Smith-Edwards et al., 2016;Chisholm et al., 2018;Guo et al., 2019). This approach allows recording of neural activities from a whole mouse DRG (Guo et al., 2019), and thus is particularly suitable for studying visceral afferents whose somata are sparsely distributed in DRGs.

To enhance the recording efficiency from visceral afferents, we here report a cost-effective imaging system that allows optical GCaMP6f recordings of DRG neurons from a wide range of thoracic, lumbar, and sacral DRGs. By using two consumer-grade cameras with two photographic lenses, we simultaneously record GCaMP6f signals at two parallel focal planes with a 1.9-by-1.3-mm field of view, a ~0.7 µm/pixel resolution, and a throughput of 50 frame per second for each camera. By tuning the ultrasonic motor ring within the photographic lenses, we can perform programmable control of axial focusing, providing a simple yet powerful tool for precise axial focus tracking without perturbing the sample (Supplementary Material 1-2). As an example, we implemented the optical recording system to characterize lumbar splanchnic and pelvic afferent innervations the colorectum. We harvested thoracolumbar (T12 to L2) and lumbosacral (L5 to S1) DRGs innervating the colorectum via the lumbar splanchnic and pelvic nerves, respectively. We evoked colorectal afferent responses by delivering mechanical stimuli to the colorectum and implemented imaging stabilization algorithms to overcome the motion artefacts from mechanical disturbance. Furthermore, we have developed and optimized the post-processing algorithms to conduct unsupervised detection of visceral afferent responses from GCaMP6f recordings (Supplementary Material 3).

## Materials and Methods

All experiments were reviewed and approved by the University of Connecticut Institutional Animal Care and Use Committee (IACUC).

### Optical recording setup

As shown in **Figure 1A-B**, the reported imaging system consists of the optical excitation and recording pathways. At the excitation path, we use a designated LED light source with narrow frequency band (470±13 nm) to excite GCaMP6f in DRG neurons. At the detection path, we use a high numeric aperture (NA) Nikon water dipping objective lens (16x, 0.8 NA) and two Canon photographic lenses (Canon EF 85mm f/1.8 USM) for image acquisition. The system offers a wide field of view (1.9 by 1.3 mm) capable to capture a whole mouse DRG. As shown in **Figure 1A**, the emitted fluorescence signals (506 – 545 nm) are evenly split into the two photographic lenses and captured by two image sensors (Sony IMX 183CLK, 2.4 µm pixel size). For both sensors, we perform 2 by 2 binning and acquire 8-bit gray-scale images at 50 frame per second. Each image resolution is 2736 by 1824 pixels with a 0.7 µm/pixel resolution at the focal plane in the DRG, sufficient to resolve the GCaMP6f signals in individual mouse DRG neurons of Φ10 - 40 µm.

**Figure 1.**
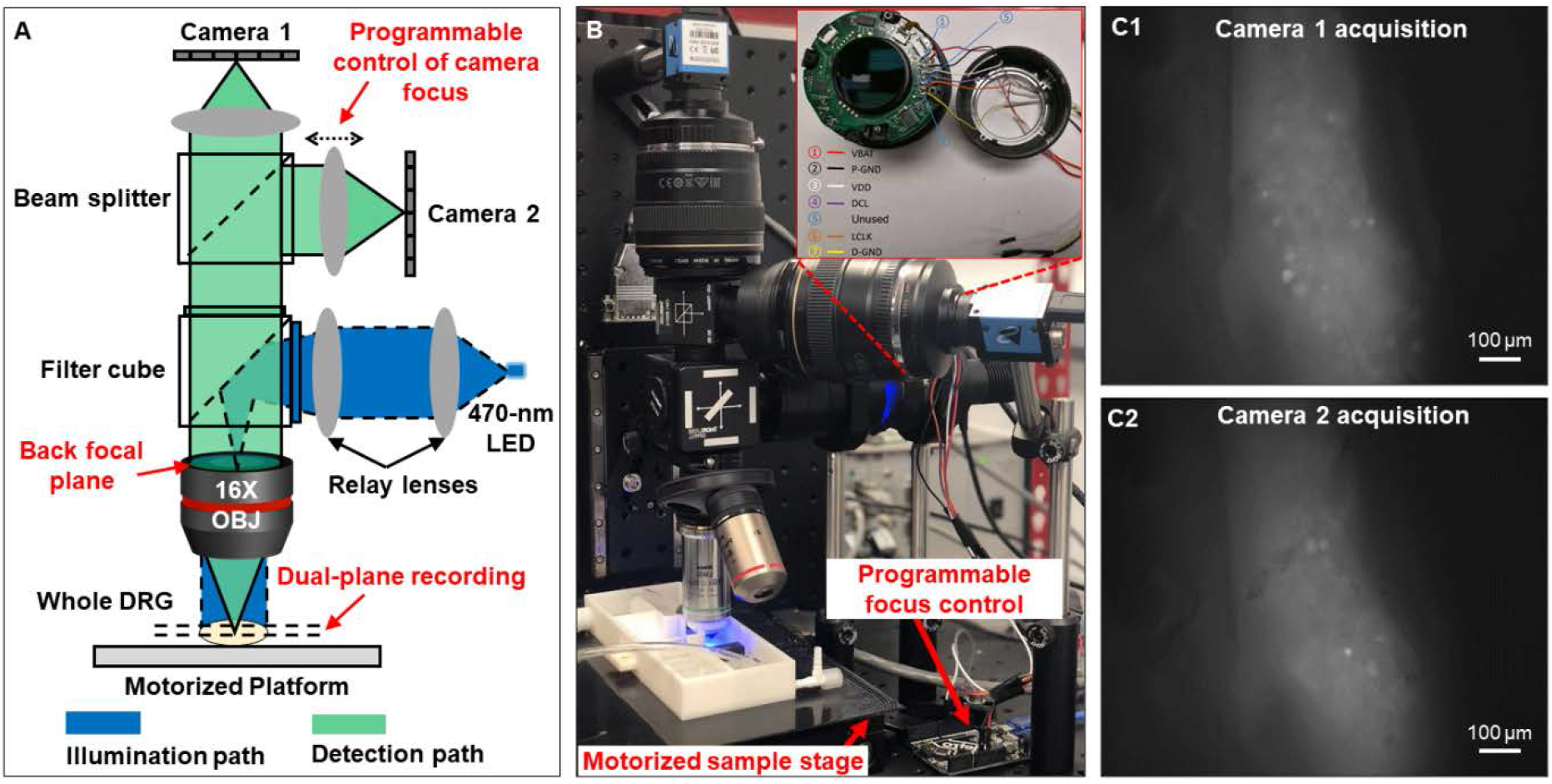
Optical setup that simultaneously captures two focal planes of a whole DRG. (**A**) The schematic of reported system. (B) The prototype setup. (C) Magnified view of the two focal planes recorded from an intact DRG.

One innovation of our microscopy system is that we can perform precise focus control using the two off-the-shelf Canon photographic lenses. For many electrophysiology experiments, axially moving the stage or the objective lens may perturb the samples, leading to image miss-alignment and movement artefacts. In our system, we perform remote focus control using the ultrasonic motor ring within the photographic lenses without perturbing the sample as shown in **Figure 1B**. The motors and control circuits inside the lens are connected to an Arduino Uno board (ATmega328P) via a 7-pin connecter. By driving the motor ring to different positions, we can adjust the focal plane by up to 160 µm with a minimum axial step of 0.1 µm. Technical details for the remote focus control are reported in Supplementary Material 2.

The motorized sample stage (MS-2000 and LX-4000, Applied Scientific Instrument) in **Figure 1B** allows maximum sample movement of 100 mm in *x* and *y* directions and 50 mm in *z* direction. The motorized platform is controlled via an open-source software, micro-manager (Edelstein et al., 2014). The stage allows imaging of biological samples across a large field of view, for example, multiple lumbosacral and thoracolumbar DRGs inside mouse vertebrae. We have also developed customized programs with graphic-user-interfaces (GUI) to allow on-screen control of the image capturing. The source codes are included in Supplementary Material 3.

### Transgenic mice

The Ai95 mice (C57BL/6 background) carrying homozygous GCaMP6f gene (strain# 28865, The Jackson Laboratory, CT) and homozygous VGLUT2-Cre mice (strain# 28863, Jackson Laboratory, CT) were crossbred. The Ai95 mice carried the gene “CAG-GCaMP6f” in the Gt(ROSA)26Sor locus, which was preceded by a LoxP-flanked STOP cassette to prevent its expression. By crossing Ai95 mice with VGLUT2-Cre mice, the Cre-expressing cell population has the STOP cassette trimmed, resulting in expression of GCaMP6f in glutamatergic neurons expressing type 2 vesicular glutamate transporter (VGLUT2), which made up the vast majority of sensory neurons innervating the colorectum (Brumovsky et al., 2011). Offspring of both sexes aged 8-14 weeks with both heterozygous GCaMP6f and VGLUT2-Cre genes (i.e., VGLUT2/GCaMP6f) were used for optical recordings.

### Ex vivo functional characterization of colorectal afferents

We implemented the above optical recording setup to characterize the afferent encoding functions by harvesting mouse colorectum, spinal nerves and ipsilateral T12 to S1 DRGs in continuity as shown in the schematic in **Figure 2A**. Mice 8-14 weeks of age were deeply anesthetized by intraperitoneal and intramuscular injection of a 0.4 mL cocktail of ketamine (120 mg/kg) and xylazine (10 mg/kg). Mice were then euthanized by perfusion from the left ventricle with modified ice-cold Krebs solution replacing sodium chloride with equal molar of sucrose (in mM: 236 Sucrose, 4.7 KCl, 25 NaHCO_3_, 1.3 NaH_2_PO_4_, 1.2 MgSO_4_·7H_2_O, 2.5 CaCl_2_, 11.1 D-Glucose, 2 butyrate, 20 acetate) bubbled with carbogen (95% O_2_, 5% CO_2_), consistent with our prior ex vivo studies on colorectal afferents (Feng and Gebhart, 2011;Feng et al., 2016). A dorsal laminectomy was performed to expose the spinal cord and the thoracolumbar and lumbosacral DRG, i.e., from T12 to S1 DRG in **Figure 2B**. The colorectum with attached DRG and vertebrae was carefully dissected via blunt dissection, and transferred to a tissue chamber superfused with 32-34 °C Krebs solution (in mM: 117.9 NaCl, 4.7 KCl, 25 NaHCO_3_, 1.3 NaH_2_PO_4_, 1.2 MgSO_4_·7H_2_O, 2.5 CaCl_2_, 11.1 D-Glucose, 2 butyrate, 20 acetate) bubbled with carbogen (95% O_2_, 5% CO_2_). The dura mater covering thoracolumbar (T12 to S2) and lumbosacral (L5 to S1) DRG was carefully removed by blunt dissection using sharp forceps (#5SF Dumont forceps, Fine Science Tools).

**Figure 2.**
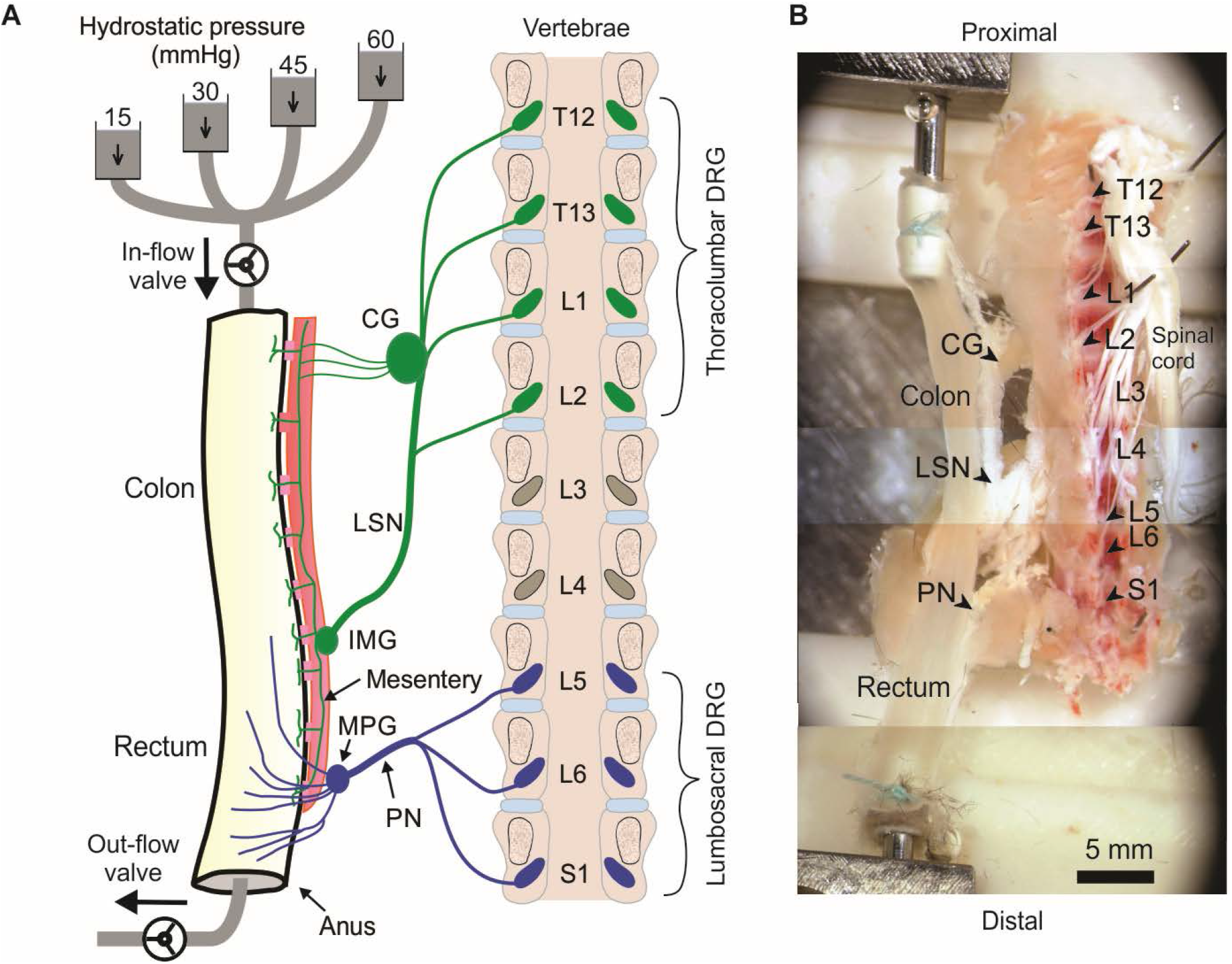
Schematic (A) and photo (B) of the ex vivo preparation for functional recordings from afferents innervating distal colon and rectum (colorectum). The colorectum, spinal nerves, and T12 – S1 DRG were harvested from mice in continuity. The colorectum was cannulated and stimulated mechanically by graded pressure distension and mucosal shearing. The photo in (B) was from stitching multiple photos taken under a stereomicroscope. LSN: lumbar splanchnic nerve; PN: pelvic nerve; IMG: inferior mesenteric ganglion; MPG: major pelvic ganglion; CG: celiac ganglion.

Since most afferents innervating hollow visceral organs are mechanosensitive (Feng and Guo, 2020), we mechanically stimulate the colorectal endings by using a custom-built colorectal distension / perfusion device as illustrated in **Figure 2A**. The colorectum was cannulated and connected to a custom-built distending device with both in-let and out-let controlled by solenoid valves. Hydrostatic pressure columns of 15, 30, 45, and 60 mmHg filled with phosphate buffered saline (PBS) were used to deliver two distinct mechanical stimuli to colorectal afferents: colorectal distension and mucosal shearing. The control function of the solenoid valves was integrated into the same MATLAB program that captures the GCaMP6f images, allowing total program-controlled mechanical stimulation and optical recording of visceral afferents. The MATLAB program controls the solenoid valves via an Arduino microcontroller. To enable the research community to easily duplicate this distending device, we have reported in details the hardware design, part information and the source code of the software in the Supplementary Materials 4.

### Optical recording of evoked fluorescence GCaMP6f signal

We captures the evoked GCaMP6f signal in each mouse DRG by high-resolution images (2736 by 1824 pixels after 2 by 2 binning), which provides a spatial resolution of 0.7 µm/pixel, sufficient to resolve individual DRG neurons. This system allows the recording of Ca^2+^ transients to resolve individual action potentials (APs) in a whole GCaMP6f-expressing DRG at two different focal planes simultaneously. The GCaMP6f signals were recorded at 50 frames per second, a sampling rate justified by the frequency spectrum of recorded Ca^2+^ transients. For a typical recording protocol of 40 sec on one DRG, a total of 4000 images are recorded, occupying 20 Giga bytes of hard drive space.

### Automated detection of GCaMP6f signals from recorded image stacks

We have developed an integrated routine to automatically extract GCaMP6f signals from recorded image stacks. The program first performs image alignment to account for the motion artefacts during DRG recording, which is unavoidable when characterizing the mechanotransduction of visceral afferents by mechanically stimulating the attached colorectum. It will then automatically detect DRG neurons with positive GCaMP6f signals using a series of unsupervised signal processing algorithms as detailed below, i.e., marker-based watershed segmentation, band-pass filtering, and variance analysis.

#### Image alignment

As illustrated in **Figure 3A**, **B1**, and **B2**, there is usually translational movement of more than 30 µm in the recorded DRG images during mechanical colorectal distension or mucosal shearing. We employ a misalignment correction algorithm to correct this translational and slight rotational movement of DRG(Mattes et al., 2001). We use the first frame as the reference image and perform image registration for all other frames in the image stack by maximizing the mutual information (MI) of different images. MI is a measure of image matching, and it does not require the signal to be the same in the two images (i.e., the second image can be slightly distorted with respect to the first one). It is a measure of how well one can predict the signal in the second image from the signal intensity in the first image. MI has been widely used to match images captured under different imaging modalities (Li, 1990;Pluim et al., 2003). The mutual information *MI* between two images *X* and *Y* can be expressed as:

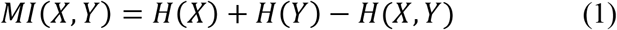
where *H*(*X*) and *H*(*Y*) are the entropy of the two images, and *H*(*X, Y*) is the joint entropy. A higher MI implies larger reduction in the uncertainty between the two distributions, which means the images are better aligned. In our implementation, we allow translational shift and image rotation in the registration process. We use a gradient decent algorithm to maximize the MI with subpixel accuracy (Van der Bom et al., 2011). To ensure the convergence, we apply 50 iterations in the optimization process. **Figure 3A** shows the first frame of the captured image stack. **Figure 3B** shows the overlays between the unaligned/aligned images of two frames. We observed a significant positional drift without applying the MI alignment process as shown in **Figure 3B1** and **B2**, which were corrected by the alignment algorithm as shown in **Figure 3B3** and **B4**. Displayed in **Figure 3C** is the quantified positional shift in x and y directions for the recorded 2000 images in one experiment, showing a maximum shift of over 40 µm. We did not plot the rotation angle as it is relatively insignificant compared to the translational shift. A representative image stack before and after alignment is converted into two videos and reported in the Supplementary Video 1.

**Figure 3.**
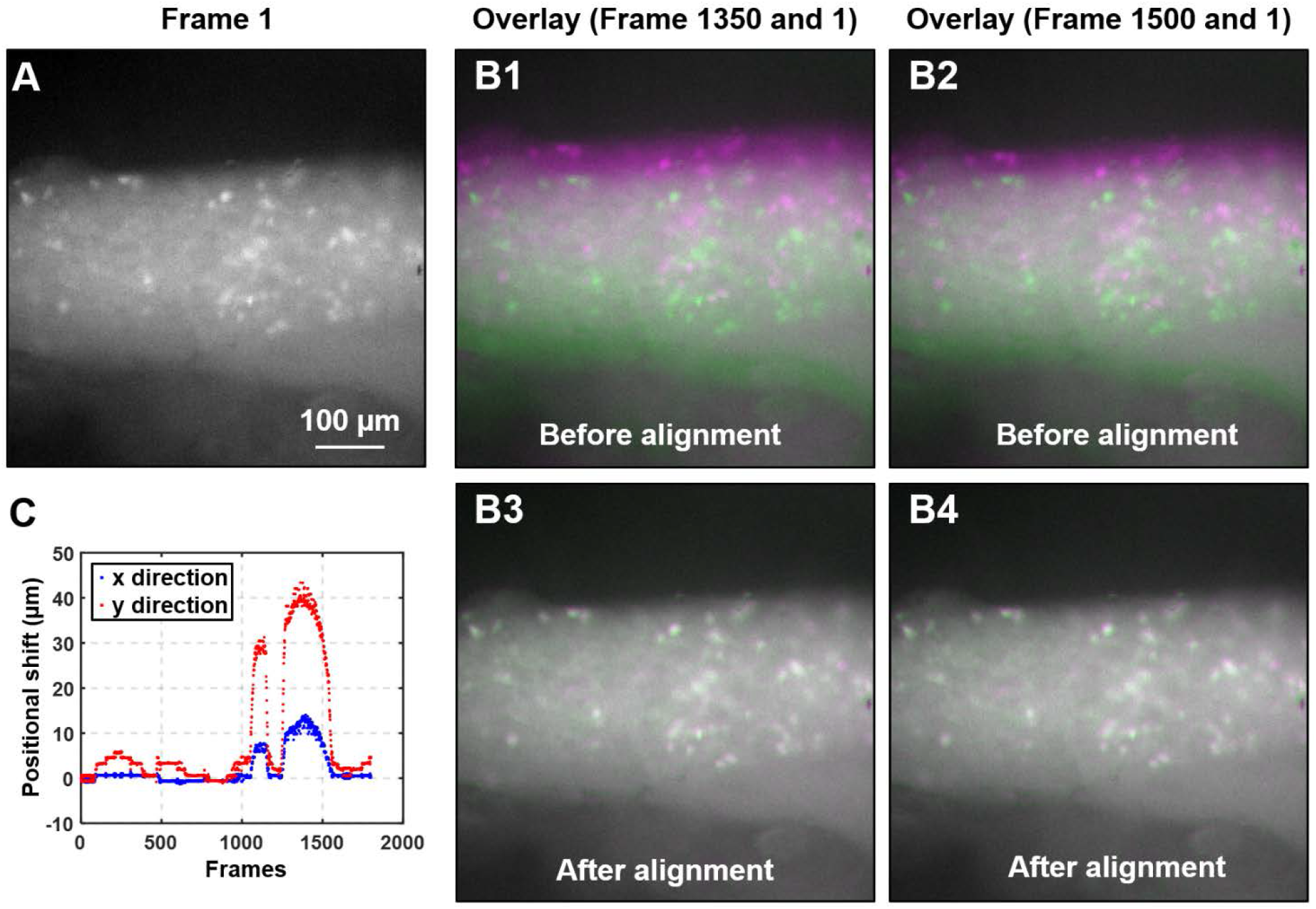
Algorithm to mitigate the motion artefact during DRG recording. (A) The magnified view of the first frame of the captured image stack. (B) The overlays of two typical image frames (#1350 and 1500, in magenta color) with the first frame (in green). Significant motion artefacts were revealed with overlays in B1 and B2 before alignment, which was greatly reduced in B3 and B4 after alignment. (C) The translational shifts of all image frames recorded from a typical colorectal distension protocol.

#### Automatic detection of GCaMP6f signals

**Table 1** summarizes the procedures of GCaMP6f signal detection. We first calculate the variance map *V* based on the aligned image stack *I_j_*(j = 1,2, ···,*J*), where *I_j_* is the *j^th^* captured image. We then initialize the global threshold *H_G_*, the estimated number of active neurons *P*, and the size range of neuron *R_min_* and *R_max_*. In the iterative neuron identification process, the variance map *V* is converted into two binary images *BW_global_* and *BW_adaptive_* using global threshold *H_G_* and adaptive threshold *H_A_*, respectively. The adaptive threshold *H_A_* is chose based on the local mean intensity in the neighborhood of each pixel. The pointwise product of the binary images *BW_global_* and *BW_adaptive_* gives a binary image *BW_combined_*, which represents the map where signals vary most at different time points. This binary map also suppresses the information outside the region of interest for better signal extraction. Next, we apply morphological closing and opening operations to *BW_combined_* as follows:

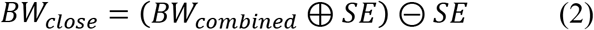

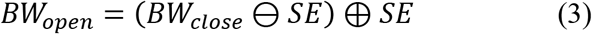
where ⊕ denotes dilation, ⊖ denotes erosion, and we use a 3-by-3 structuring element *SE* of [0,1,0;1,1,1;0,1,0] in Eqs. (2)–(3). These two morphological operations can clean up the background noise for better signal extraction. We further remove the small features that have fewer than 40 pixels in size from the binary result *BW_open_* and clear the features at image borders. The updated binary result *BW_updated_* is then used to create the distance matrix *M_distance_* as follows:

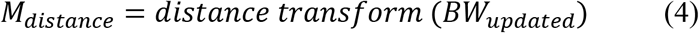
where the distance transform of a binary image is the distance from every pixel to the nearest nonzero-valued pixel of that image. The watershed transform of the distance matrix *M_distance_* then returns a label matrix *L_watershed_* that identifies the possible active neurons’ locations (watershed regions). The predefined neuron size range *R_min_* and *R_max_* are used to select the expected neurons and update the label matrix as *L_selected_*. At the end of this iterative process, the global threshold *H_G_* is updated by a step size *α*. In **Figure 4B**, we plot the GCaMP6f signals of identified neurons for further analysis.

**Figure 4.**
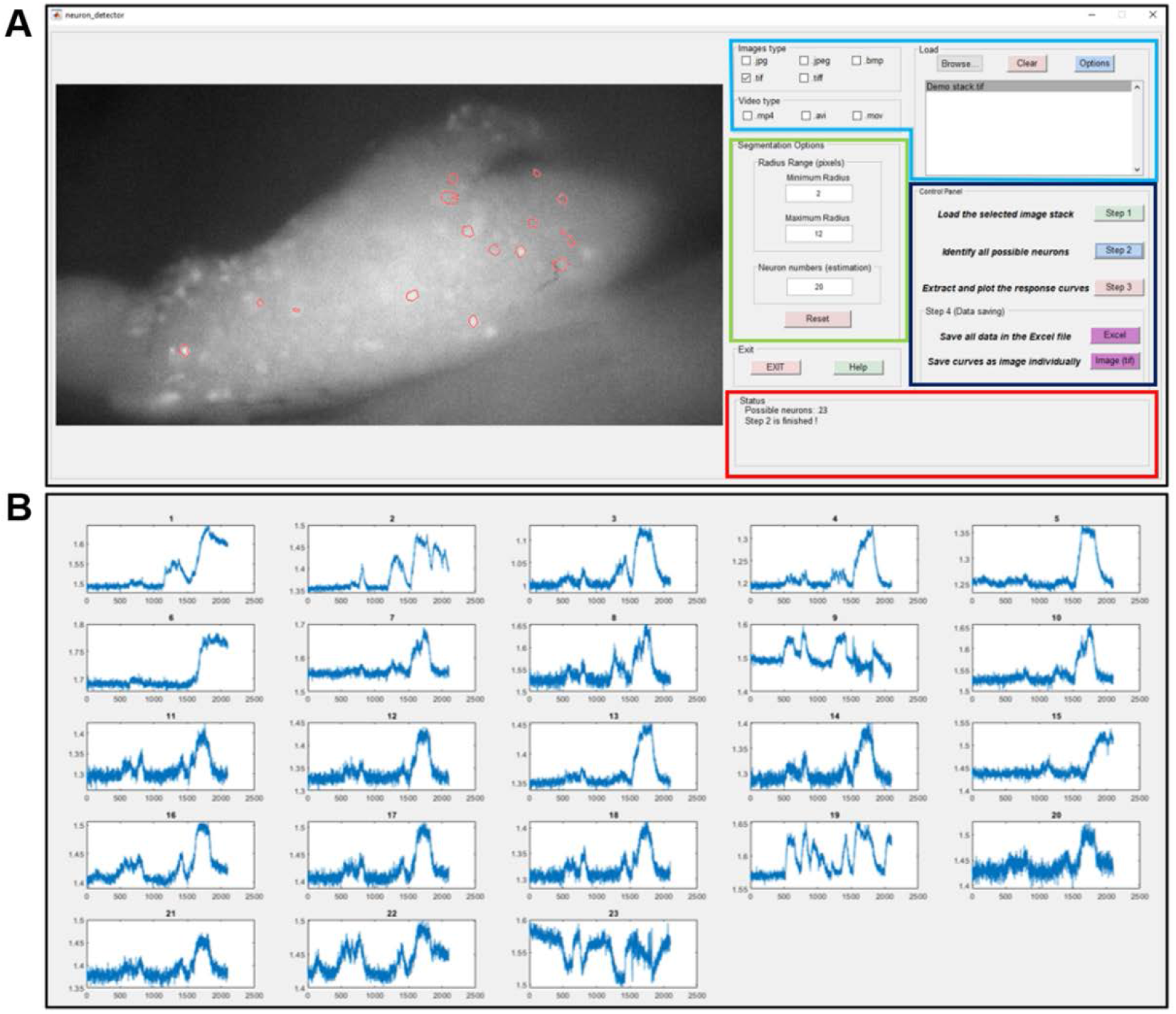
The graphic-user-interface (GUI) of the custom-built software for automatic extraction of GCaMP6f responses from image stacks. (A) The GUI with five panels: The image process window (right), the input image format selection panel (right top with blue label), segmentation options (right middle with green label), control panel (right middle with black label) and status panel (right bottom with red label). (B) The extracted intensity profiles of evoked DRG neurons. Detailed descriptions are listed in Supplementary Material 3.

**Table 1.**
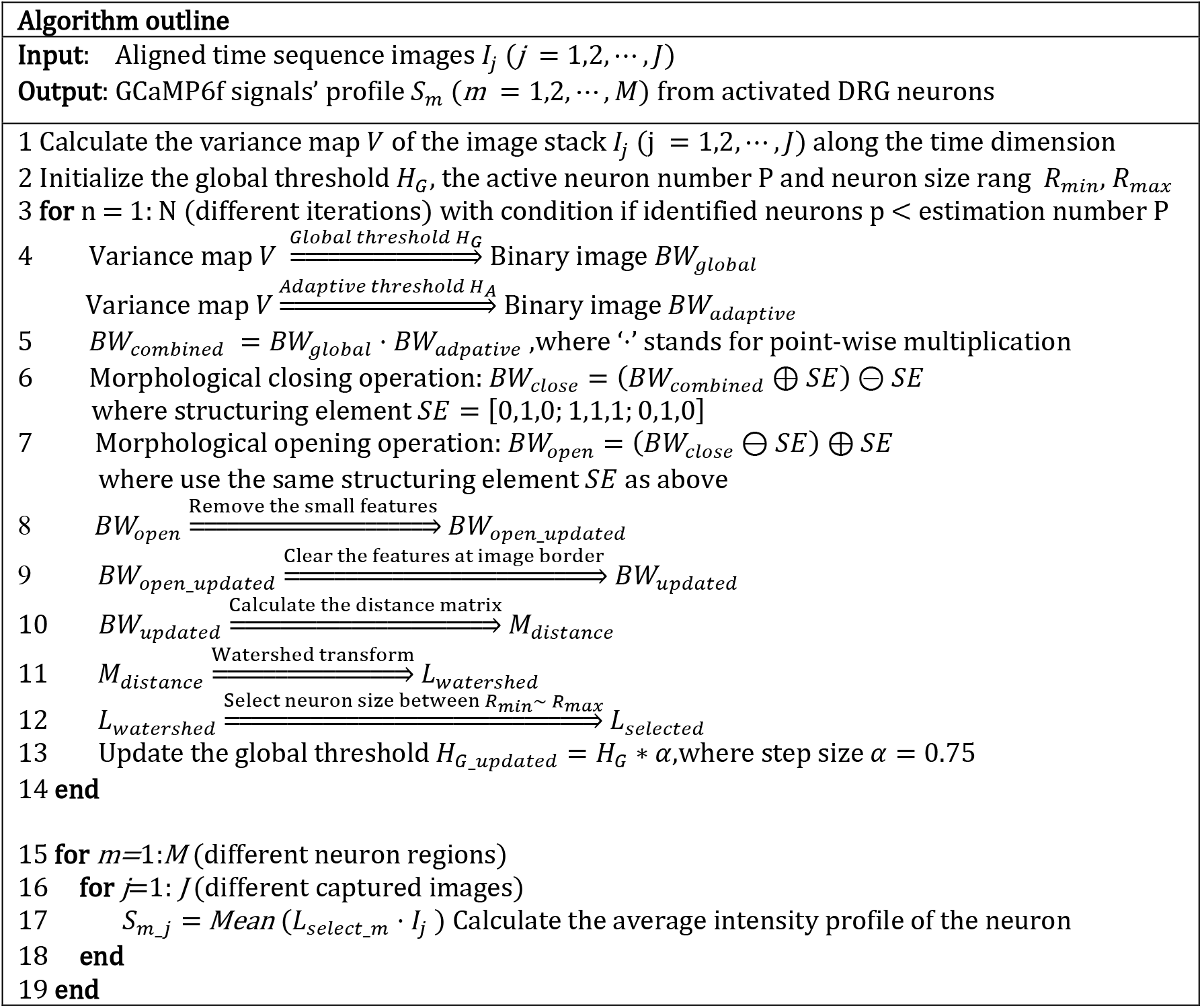
Algorithm outline for automatic detection of GCaMP6f signals

### Afferent identification and classification

Mouse colorectal afferents were activated by two physiologically correlated stimuli at the colorectum: stepped luminal distension by hydrostatic fluid column of phosphate buffered saline (PBS, 15, 30, 45, and 60 mmHg of 5-sec steps) and luminal shear flow of PBS (20-30 mL/min) (Guo et al., 2019). Based upon response profiles to graded distension and luminal shear, colorectal DRG were functionally classified into four classes: low-threshold (LT) muscular, high-threshold (HT) muscular, mucosal, and muscular-mucosal classes. LT-muscular afferents responded to all four distension pressure levels whereas HT-muscular only responded to noxious distension pressure (30, 45 and 60 mmHg); colorectal intraluminal pressure beyond 20 mmHg was considered noxious to mice (Kamp et al., 2003;Feng et al., 2010). Mucosal afferents did not respond to distension but responded to luminal shear flow. Muscular-mucosal afferents responded to both luminal shear flow and colorectal distension at all four pressure levels.

### Data recording and analysis

Extracted GCaMP6f signals in the form of pixel intensity (0-255) from individual DRG neurons were normalized by the pre-stimulus intensity. Peak GCaMP6f transients were determined when the signal increased by 3% within 200 mSec. The duration of the GCaMP6f transients were determined by the measuring temporal width of the signal at 25% of the peak intensity. Proportions of afferent classes were compared by Chi-square test using SigmaStat v4.0 (Systat software, Inc., San Jose, CA). P < 0.05 was considered significant.

## Results

Using our custom-built imaging system, the evoked GCaMP6f transient in multiple DRGs were recorded at individual neural resolution. Displayed in **Figure 5** are evoked GCaMP6f transients (ΔF/F, normalized fluorescent signals) in DRG neurons by electrical stimulation of attached dorsal roots. Recordings were conducted at 0.5, 2 and 4 Hz stimulation frequency (**Figure 5A**). The evoked GCaMP6f transients in **Figure 5B** showed unanimous increase in baseline GCaMP6f intensity when stimulus frequency is beyond 0.5 Hz. The duration of the GCaMP6f transients were measured from 11 neurons at 0.5 Hz stimulation, showing an average duration of 1.31 ± 0.19 sec. Displayed in **Figure 5C** are the frequency domain of the GCaMP6f transients via fast Fourier transform, which showed that the majority of the signal frequency is between 0.3 and 5 Hz. The GCaMP6f signals were band-pass filtered (0.3 to 5 Hz) and displayed in **Figure 5D**. By analyzing the filtered signal, evoked single-spike GCaMP6f transients can be reliably detected in all recordings from 16 DRG neurons at 0.5 and 2 Hz stimulation, showing peak-to-peak ΔF/F above 3.5%. At 4-Hz stimulation, only 15% of the recordings (13 out of 87 DRG neurons) allow reliable detection of single-spike GCaMP6f transients (peak-to-peak ΔF/F above 3.5%) whereas the others do not.

**Figure 5.**
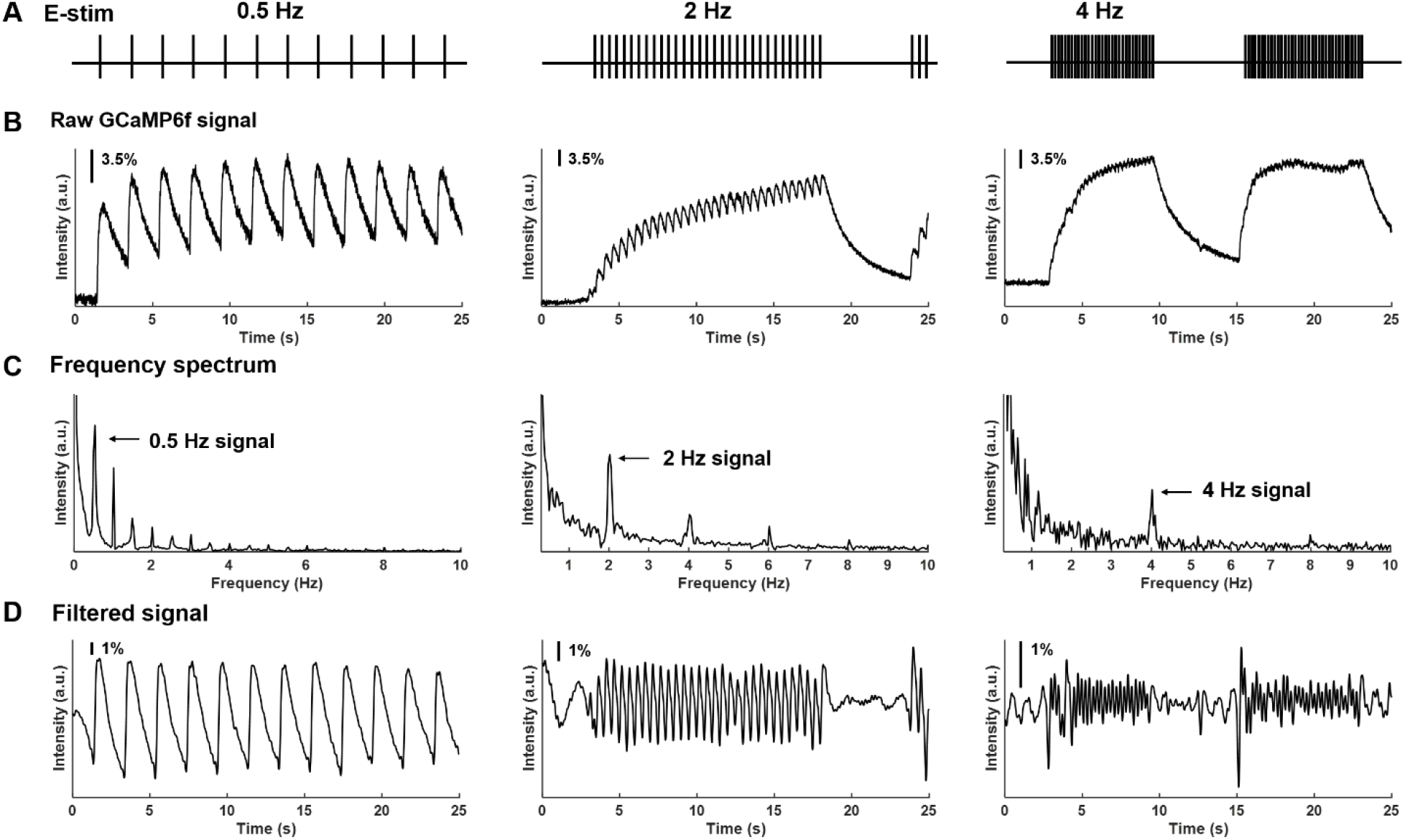
Frequency spectrum analysis on GCaMP6f responses evoked by electrical stimulation of the attached dorsal root. (A) The implemented electrical stimulation at 0.5, 2 and 4 Hz, respectively. (B) Evoked GCaMP6f signals in DRG neurons. (C) The frequency domain of the GCaMP6f transients from fast Fourier transform. (D) Band-pass filtered GCaMP6f signals.

In addition to electrical stimulation, mechanical colorectal distension and mucosal shearing were implemented to evoke GCaMP6f transients in colorectal DRG neurons (**Figure 6A-B**). The frequency domain of the GCaMP6f transients in **Figure 6C** indicates that the 0 to 5 Hz range covers most, if not all the signals of the GCaMP6f transients. Evoked afferent spikes by colorectal distension are generally high frequency (> 2Hz) to prevent reliable detection of single-spike GCaMP6f transients. In contrast, evoked afferent spikes by mucosal shearing are usually below 2 Hz and can be detected with single-spike resolution. The colorectal GCaMP6f signals were either low pass filtered (0 to 5 Hz for colorectal distension) or band-pass filtered (0.3 to 5 Hz for mucosal shearing) and displayed in **Figure 6D**.

**Figure 6.**
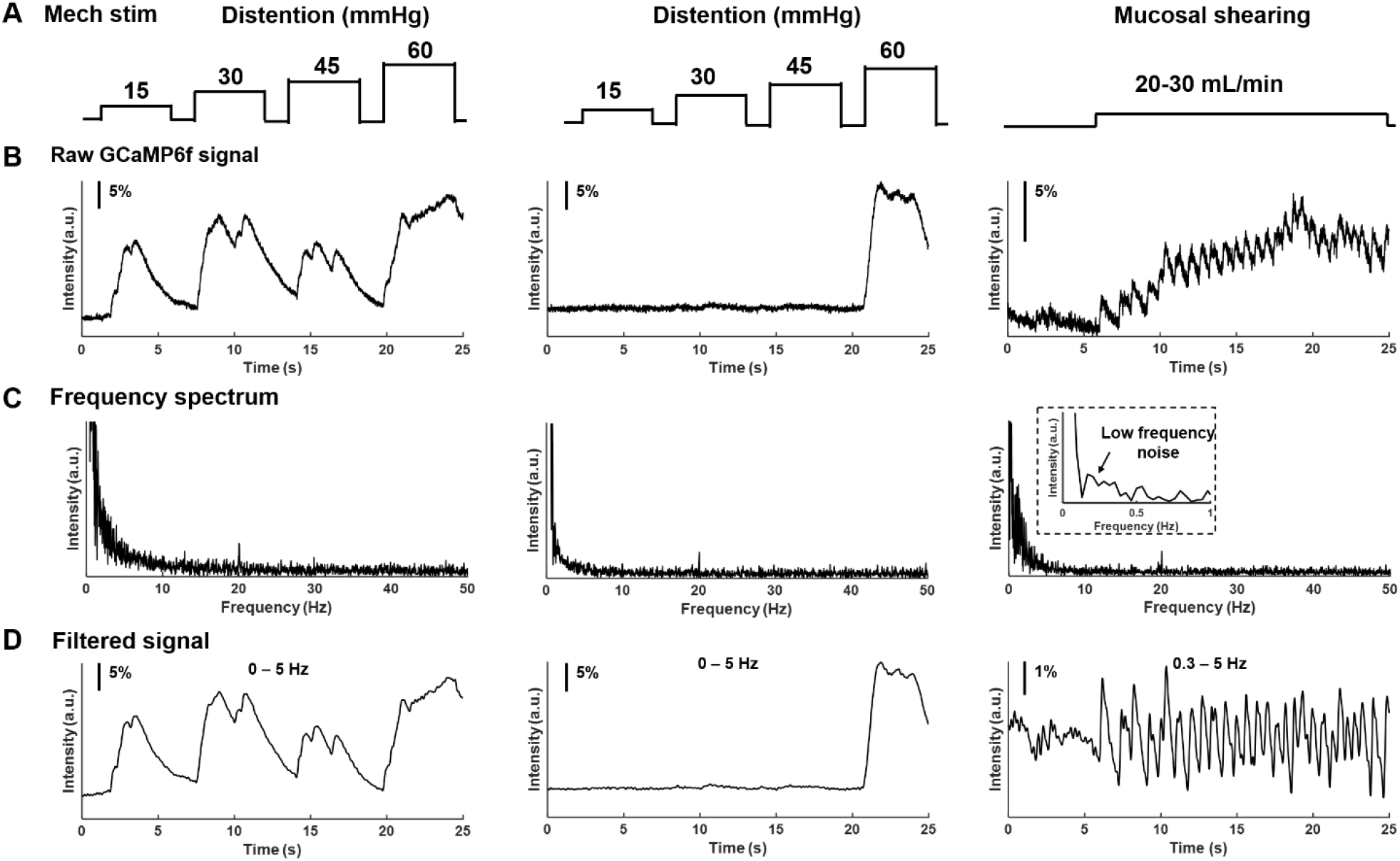
Frequency spectrum analysis on GCaMP6f responses from colorectal afferents evoked by mechanical colorectal distension and mucosal shearing. (A) The implemented mechanical colorectal distension and luminal shear flow, respectively (B) Evoked GCaMP6f signals in DRG neurons. (C) The frequency domain of the GCaMP6f transients from fast Fourier transform. (D) Band-pass filtered GCaMP6f signals.

Using our high-throughput imaging system, we recorded a total of 456 colorectal neurons that respond to mechanical colorectal distension and/or mucosal shearing. As shown in **Figure 7A**, 38.8% of the colorectal neurons are in the thoracolumbar DRG (T12 to L2) whereas 61.2% in the lumbosacral DRG (L5 to S1). Colorectal afferent neurons were functionally classified into four groups based upon their response profiles to colorectal distension and/or mucosal shearing as illustrated in **Figure 7B**. The proportion of functionally distinct afferent groups in both thoracolumbar and lumbosacral innervation pathways were displayed in **Figure 7C**.

**Figure 7.**
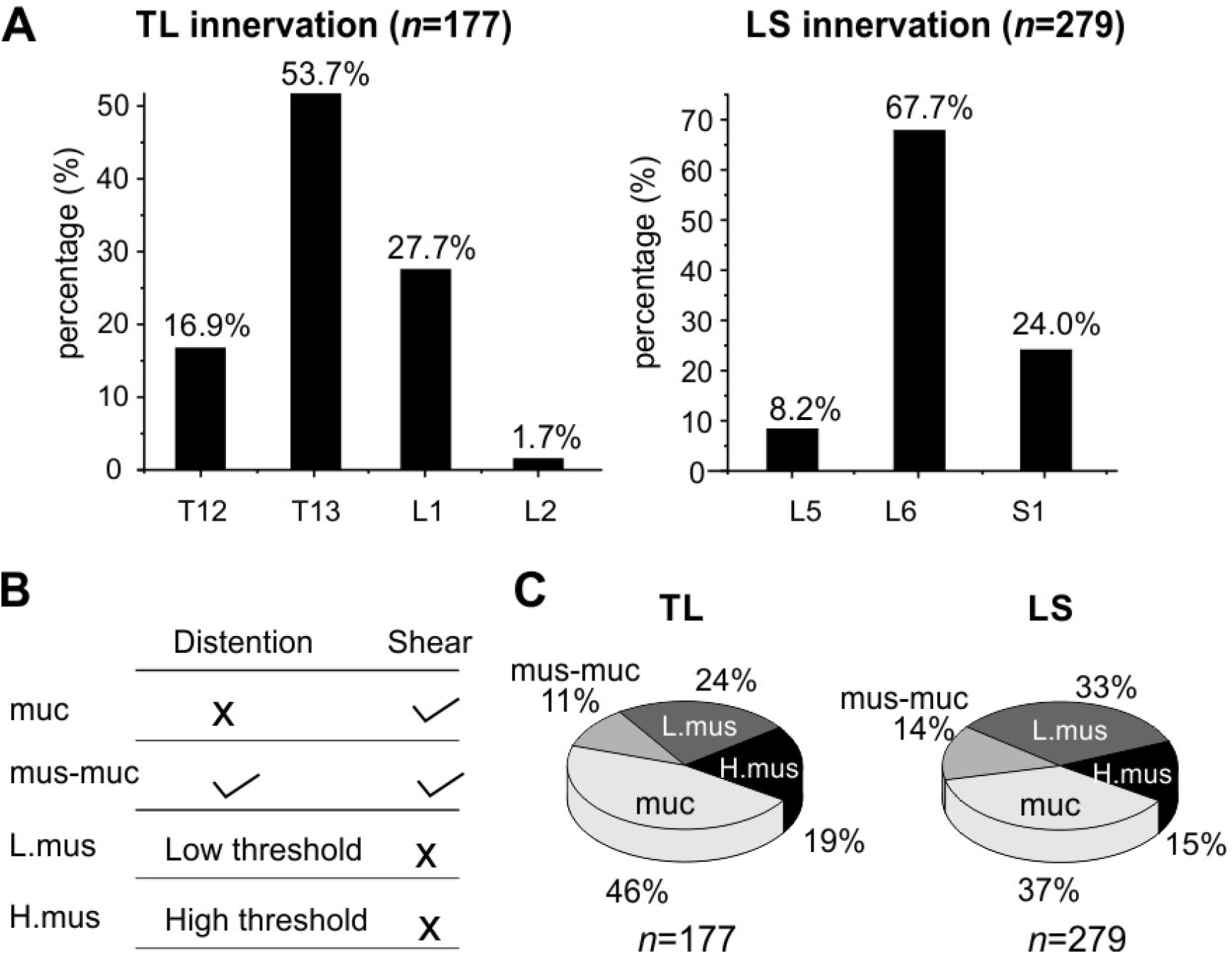
Functional recording and characterization of afferents innervating mouse colorectum in both thoracolumbar (TL) and lumbosacral (LS) pathways. (A) The distribution of colorectal neurons in the thoracolumbar (T12 to L2)) and lumbosacral (L5 to S1) DRG. (B) Functional classification of colorectal afferents based upon response profiles to colorectal distension (15, 30, 45, 60 mmHg) and mucosal shearing. (C) The distributions of four colorectal afferents classes within TL and LS innervations. muc: mucosal afferents; mus-mu: muscular-mucosal afferents; L.mus: low-threshold muscular afferents; H.mus: high-threshold muscular afferents.

## Discussion

In this study, we reported a cost-effective high-throughput approach for functional characterization of afferents innervating visceral organs, which are generally challenging by conventional electrophysiological recordings due to the sparse nature of visceral innervations. Functional characterization of neurons by optical recordings via genetically encoded calcium indicators (Podor et al., 2015) are routinely conducted in the central nervous system (CNS) where stimulation modalities are usually either electrical or chemical. Peripheral sensory neurons encode additional stimulus modalities that are generally absent in the CNS, e.g., thermal and mechanical stimulations. Functional characterization of peripheral afferents requires applying stimuli to their nerve endings embedded in the end organs of innervation. Mechanical stimulation poses the greatest challenge for optical recordings compared to other stimuli (electrical, thermal, and chemical) due to the unavoidable motion artefacts of samples during mechanical disturbance. Even slight movement of tens of microns will lead to false optical recordings from neural somata the diameter of which are usually in the same order.

Mechanical neural encoding is particularly crucial for visceral sensation and nociception (see (Feng and Guo, 2020) for a recent review). The dominant perceptions from the viscera are discomfort and pain that are reliably evoked by mechanical distension of hollow visceral organs, but not by other pain-evoking stimuli to the skin like pinching, burning, inflammation, and cutting (Feng and Guo, 2020). Despite the importance of visceral mechanotransduction, there has been only one report in the literature characterizing mechanical visceral neural encoding by optical recordings (Guo et al., 2019), largely due to the challenge of movement artefacts. Another major challenge for conducting optical neural recordings is the high cost of the optical setups, the most widely used of which are two-photon and confocal scanning fluorescent microscopes. This prohibits a wider application of optical recordings in studying visceral afferents. We recently implemented a scientific CMOS camera with 82% quantum efficiency (Zyla-4.2P, Andor Technology, South Windsor) and demonstrated feasibility to conduct full-frame recordings of GCaMP6f signals in the DRG by epi-fluorescent excitation at 100 frames per second (Guo et al., 2019). However, the cost of those scientific CMOS cameras is not trivial. In current study, we successfully addressed the above limitations in our custom-built optical setup by 1) applying a robust image alignment algorithm to account for the translational and rotational movement of neural samples, 2) using consumer-grade optical components and image sensors to assemble the whole setup from scratch within a limited budget, and 3) further reducing sample perturbation via remote focusing using the ultrasonic motor rings of the photographic lenses. We foresee a wider adoption of this approach by the research community, which will likely expedite the functional characterization of visceral afferents as well as neurons innervating non-visceral organs.

Compared with sensory innervations of the extremities, the DRGs are in closer proximity with visceral organs and thus more susceptible to mechanical stimuli to their nerve endings in the organ wall. In current study, the recorded DRG images during mechanical colorectal distension of 60 mmHg can undergo translational movement of up to 50 microns and rotational movements of about 2 degrees. This has confounded the extraction of GCaMP6f signals from mouse colorectal DRG neurons which are generally 10 – 40 microns in diameter (Christianson et al., 2006a;Christianson et al., 2006b). To the best of our knowledge, this is the first report to document the application of an alignment algorithm to compensate image recordings of the DRG. Compared with generic alignment algorithm based upon cross correlation analysis (e.g., (Guizar-Sicairos et al., 2008)), we implemented an MI-based algorithm that do not require images to be identical (DRGs indeed showed different GCaMP6f intensity and patten during mechanical stimulation protocols). After the alignment process, contours of individual DRG neurons generally fall within a margin around 3 microns wide, sufficiently small to avoid interfering the ensuing extraction of GCaMP6f signals from individual somata.

We have further reduced the cost by assembling the optical recording setup using consumer-grade components, including the Canon photographic lenses and the SONY image sensors (IMX 183CLK). Compared with high-cost scientific CMOS cameras, the slightly lower sensitivity and quantum efficiency of the SONY sensor require about twice the exposure time as the scientific one to achieve comparable imaging quality. We conducted frequency spectrum analysis of recorded GCaMP6f transients and demonstrated that 0 to 5 Hz is the dominant frequency range of the evoked colorectal afferent activities, consistent with the maximum spike frequencies of about 5 Hz in mouse colorectal afferents from prior electrophysiological studies (Feng et al., 2010;Feng et al., 2012a). In addition, the relative low cost of the SONY sensor allows us to adopt two cameras to simultaneously record at two parallel focal planes, i.e., recording from a volume of DRG tissue to double the efficiency. Instead of using a conventional microscope tube lens, we employ two Canon 85-mm photographic lenses in our platform. The photographic lens allows us to perform remote axial focus control with high spatial precision of 0.1 µm. By tuning the ultrasonic motor ring to different positions, the evoked GCaMP6f signals at the different planes of the DRG can be recorded without perturbing the samples. Also, using commercial grade photographic lens for remote focus control significantly reduces the cost of our imaging system compared with high-cost piezo stages used in conventional microscopes.

We also implemented an unsupervised algorithm to allow automatic extraction of DRG neurons with positive GCaMP6f responses from recorded image stacks. The major advantage is that it can be used for various fluorescence imaging systems with different research purposes. By adjusting the estimated neuron size and number options in the GUI, users can modify the routine for their experiment systems. Both image format (like jpeg, tiff, and bmp) and video format data (like mp4, avi, and mov) can be processed using the reported GUI. We note that the recording and data processing is also not demanding in computing power, only requiring a personal computer with a modern CPU, 32 GB RAM and a solid state drive. Compared with the usual manual process of marking neurons, our procedure allows expedited and unbiased processing of large amount of image data in a robust and reliable fashion. We anticipate the adoption of this processing routine in the neuroscience research community for increasing the efficiency of extracting neural responses from large datasets.

The calcium indicator GCaMP6f produces stronger fluorescent signals than GCaMP3 and 5 and have faster recovery kinetics than GCaMP6m, 6s, 7(Ohkura et al., 2012;Chen et al., 2013;Muto et al., 2013), making it ideal for characterizing single-spike neural activities (Podor et al., 2015). We measured the temporal width of the GCaMP6f transients of individual action potential spikes in mouse DRG to be close to 1.3 sec, indicating complete recovery to baseline Ca^2+^ fluorescent levels when the spike frequency is below 0.5 Hz. We showed in current study that spiking frequency below 2 Hz can be reliably determined in single-spike resolution from GCaMP6f transients in all DRG neurons whereas frequency at 4 Hz can be determined in only 15% of the DRG neurons. This discrepancy in determining single spikes at 4 Hz stimulation likely reflects the different intracellular Ca^2+^ events in different DRG neurons. Consistent with the prior findings (Chisholm et al., 2018;Hartung and Gold, 2020), spike frequencies beyond 4 Hz will generally results in a large GCaMP6f transient, from which single spikes usually cannot be determined. A recent systematic study on mouse trigeminal ganglion neurons indicates that the magnitude and rate of those large GCaMP6f transients are not reliable measures of neural activity, nor can be used to assess changes in activities (Hartung and Gold, 2020). Hence in current study, we used GCaMP6f responses to exclusively assess whether visceral neurons responded to certain mechanical stimuli to the colorectum and use their response profiles to functionally separate them into different groups. Our optical approach allows functional characterization of 456 afferents innervating the colorectum, reporting a large number of afferents than previous approaches using single-fiber electrophysiological recordings (Brierley et al., 2004;Feng and Gebhart, 2011). To identify colorectal afferents in the thoracolumbar (TL) and lumbosacral (LS) pathways, single-fiber recordings were conducted on manually teased fine nerve filaments from the lumbar splanchnic (LSN) and pelvic nerves (PN), an approach that will not determine the relative innervation densities between the two pathways. In current study, this non-biased optical recording approach allows us to determine that thoracolumbar pathway makes up a much smaller proportion of the total afferent innervation (39%) than the lumbosacral pathway (61%). Within the LS pathway, the proportion of mechanosensitive afferents is comparable to our previous report (Guo et al., 2019).

Interestingly, mucosal afferents that encode luminal shearing makes up a significant proportion in the TL pathway from current study, which contrasts with the limited number of mucosal afferents characterized by single-fiber recordings from the LSN (Brierley et al., 2004;Feng and Gebhart, 2011). We speculate that this is due to the slightly stronger mechanical stimuli of mucosal shearing induced by fluid flow in the tubular colorectum in current study than the fine mechanical stroking by a 10 mg fine brush on flattened colorectum in previous studies (Brierley et al., 2004;Feng and Gebhart, 2011). In addition, the ex vivo preparation in current study also includes small fibers that by-pass the celiac ganglion, an innervation pathway that was absent in previous studies when recordings were conducted exclusively from the LSN distal to the celiac ganglion. Further research is warranted to identify the exact innervation pathways that contribute to the neural encoding of colorectal mucosal shearing by thoracolumbar DRG, likely via nerve transection studies.

## Conclusion

In conclusion, we report a turnkey microscopy system that allows Ca^2+^ imaging of DRG neurons from a wide range of thoracic, lumbar, and sacral DRGs. By using two consumer-grade image sensors with two photographic lenses, we simultaneously record GCaMP6f signals at two parallel focal planes with a 1.9-by-1.3-mm field of view, a ~0.7 µm/pixel resolution, and a throughput of 50 frame per second from each camera. By tuning the ultrasonic motor ring within the photographic lenses, we achieved programmable focus control in axial direction, providing a simple yet powerful tool for precise axial focus tracking without perturbing the sample. The custom-built post processing software implemented an image alignment algorithm based on mutual information to address motion artefacts and achieved automatic extraction of the GCaMP6f signals from DRG image stacks with high computing efficiency. As a demonstration, we functionally characterized 456 afferents innervating mouse colorectum in both thoracolumbar and lumbosacral pathways. The reported system offers a cost-effective solution for recording visceral afferent activities from a large volume of DRG tissues.

## Data Availability Statement

All datasets generated for this study are included in the article/supplementary material.

## Ethics Statement

All experiments were reviewed and approved by the University of Connecticut Institutional Animal Care and Use Committee

## Author Contributions

Z. B., T. G. and S. J. prepared the display items. B. F. prepared the initial draft of the manuscript. All authors contributed to all aspects of manuscript preparation, revision, and editing. B. F. and G. Z. supervised the project.

## Acknowledgments

This work was supported by NINDS U01 NS113873 awarded to Drs. Feng and Zheng and NIDDK R01 DK120824 grants awarded to Dr. Feng.

