## Supplementary material 1 for "High-throughput functional characterization of visceral afferents by optical recordings from thoracolumbar and lumbosacral dorsal root ganglions"

### Part list and estimated cost for the system

| Item | Specification | Amount | Cost (U.S. dollars) | Vendor or supplier |
|---|---|---|---|---|
| Aluminum Breadboard | MB18 18" x 18" x 1/2" | 1 | $281.56 | Thorlabs |
| Aluminum Breadboard | MB1218 12" x 18" x 1/2" | 1 | $193.64 | Thorlabs |
| Mounted Led | M470L4 470 nm,760mW | 1 | $296.50 | Thorlabs |
| Collimation Adapter | COP5-A | 1 | $236.98 | Thorlabs |
| Double Convex Lens | LB1630-A Ø2", f=100mm | 1 | $45.72 | Thorlabs |
| Kinematic Filter Cube | DFM1L | 1 | $386.17 | Thorlabs |
| GFP Filter Set | MDF-GFP GFP Excitation, Emission, and Dichroic Filters | 1 | $690.39 | Thorlabs |
| Beamsplitter Cube | CCM1-BS013 30 mm Cage Cube with Beamsplitter | 1 | $296.50 | Thorlabs |
| Objective Lens Turret | OT1 | 1 | $340.87 | Thorlabs |
| Lens Tube | SM1L03 | 2 | $12.52 | Thorlabs |
| Lens Tube | SM1L05 | 1 | $12.97 | Thorlabs |
| Lens Tube | SM1L15 | 1 | $16.17 | Thorlabs |
| Lens Tube | SM1M05 | 1 | $13.32 | Thorlabs |
| Lens Tube | SM2L03 | 1 | $24.06 | Thorlabs |
| Lens Tube | SM2L10 | 1 | $30.99 | Thorlabs |
| Lens Tube | SM2L15 | 1 | $31.79 | Thorlabs |
| Thread Adapter | SM1A2 | 2 | $26.51 | Thorlabs |
| Thread Adapter | SM1A4 | 1 | $24.43 | Thorlabs |
| Thread Adapter | SM2A6 | 1 | $26.51 | Thorlabs |
| Thread Adapter | RMSA10 | 1 | $24.35 | Thorlabs |
| Lens Tube | SM1RC | 1 | $25.10 | Thorlabs |

| | | | | |
|---|---|---|---|---|
| Slip Ring | | | | |
| Optical Post | TR1<br>Ø1/2", L = 1" | 4 | $4.88 | Thorlabs |
| Optical Post | TR4<br>Ø1/2", L = 4" | 3 | $6.05 | Thorlabs |
| Optical Post | TR6<br>Ø1/2", L = 6" | 3 | $7.33 | Thorlabs |
| Post Holders | UPH1<br>Ø1/2", L = 1" | 4 | $32.20 | Thorlabs |
| Post Holders | UPH3<br>Ø1/2", L = 3" | 4 | $33.83 | Thorlabs |
| Lens Tube Slip Rings | SM2RC<br>Ø2.20" | 2 | $30.99 | Thorlabs |
| Slotted Base | BA4 | 2 | $40.31 | Thorlabs |
| Right-Angle Brackets | AB90A | 3 | $27.85 | Thorlabs |
| Angle Post Clamps | SWC | 1 | $25.00 | Thorlabs |
| Lens adapter | 58mm to 52mm Step-Down Ring | 1 | $12.90 | Amazon |
| XYZ Automated stage | ASI MS2000 Stage<br>ASI LX-50A Stage<br>Control Box | 1 | $ 1,700 (used) | eBay |
| Objective Lens | Nikon Plan Fluorite Water Dipping Objective 16x 0.8 NA, | 1 | $5,695 | Edmund Optics |
| Image sensor | DMK 33UX183 | 2 | $700.0 | The Imaging Source |
| Camera Lens | Canon EF 85mm f/1.8 USM | 2 | $419.0 | Amazon |
| | | **Total** | **$13,297** | |
