## Supplementary material 2 for "High-throughput functional characterization of visceral afferents by optical recordings from thoracolumbar and lumbosacral dorsal root ganglions"

### Remote focus control using the ultrasonic motor rings of the Canon photographic lenses

#### Step 1: Parts and Tools Preparation [1]

Below are the tools for lens modification.

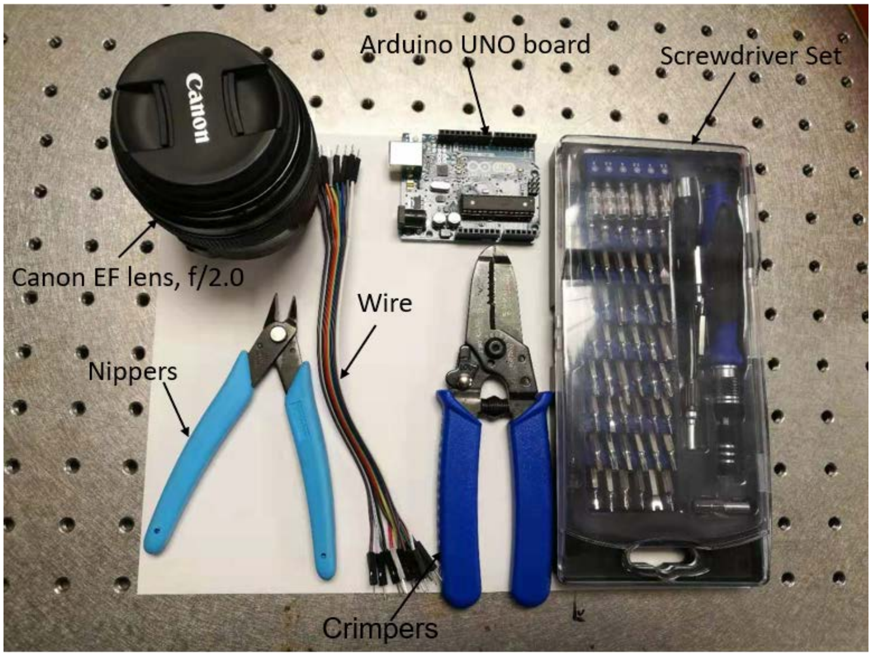


Figure 1: Tools preparation.

#### Step 2: Lens Modification [1]

- Remove the four screws on the back side of the lens.

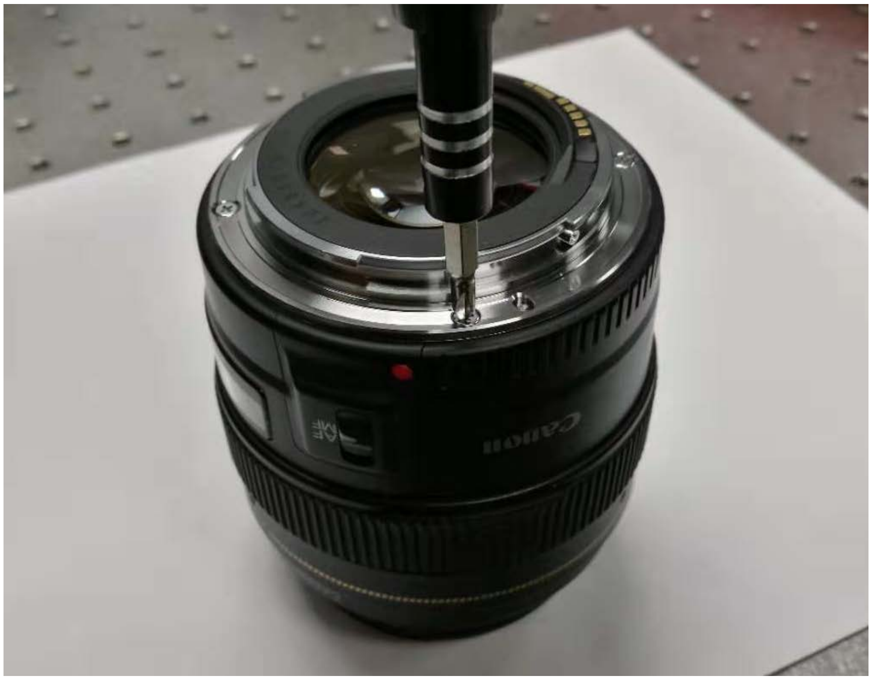


Figure 2: Remove the four screws.

- Detach the back cover and solder the wires.

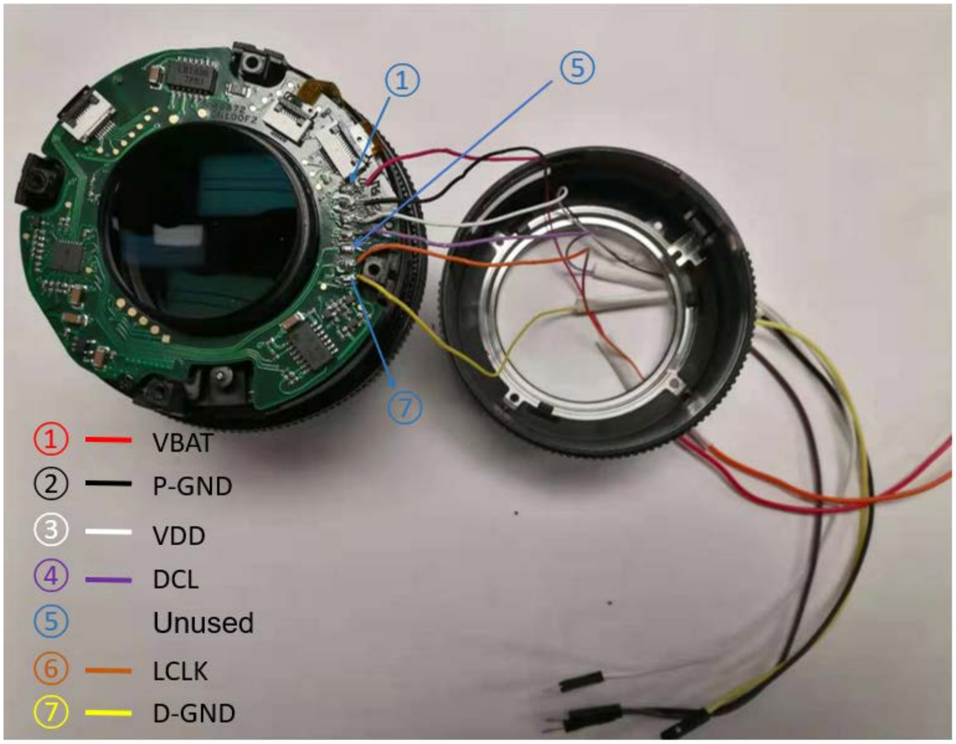


Figure 3 Detach the back cover and solder the wires.

- Reassemble

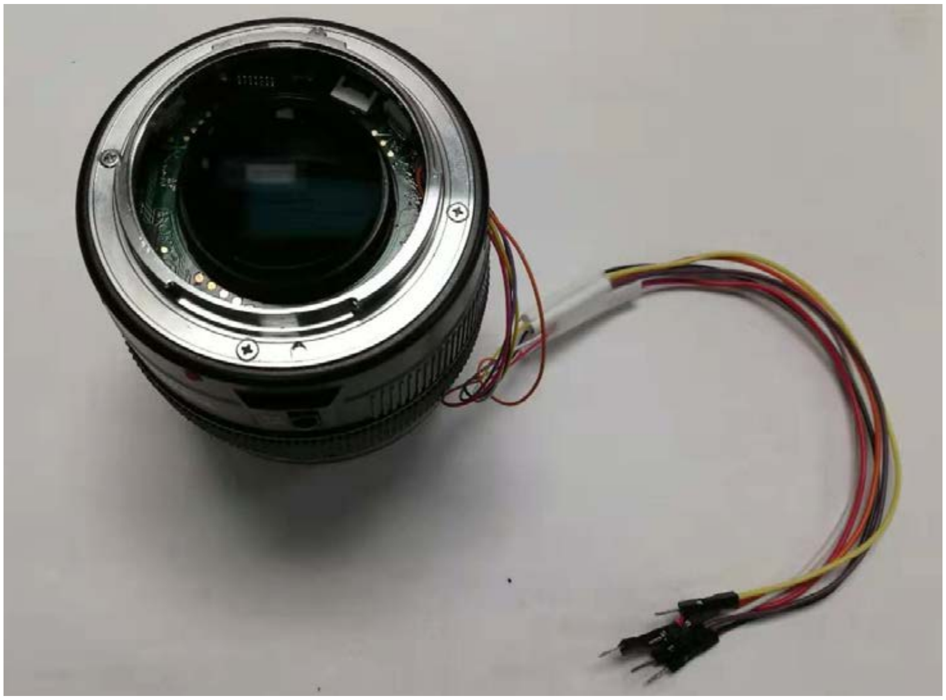

Figure 4: Reassemble the lens parts.

**Step 3: Wire Connection [1]**

The lens has motors and control circuits inside and can be connected to an external board via a 7-pin connector. Three pins for the power supply and four pins for a standard SPI interface. We need to wire the pins from outside so that we can send commands from an external board to move the lens ring to the in-focus position. We use an Arduino Uno board (ATmega328P) to perform focus control.

Pin 11 below is defined to send the ‘moving’ command and is connected to the DCL pin of the lens. Pin 13 is used for transferring the clock, which is connected to the CLK pin on the lens. Two GND pins are connected to the D-GND pin and P-GND pin on the lens, respectively. One 5V pin gives power supply for the lens control circuits and is connected to the VDD pin on the lens. We define an output pin as the digital logic high for the power supply of lens motors. The pinout diagram and schematic are listed below.

| **Digital logic** | | |
|---|---|---|
| **Controller** | **Lens** | **Description** |
| 5V | VDD | Power supply for digital circuit in lens |
| GND | D-GND | VDD ground |
| MOSI (pin 11) | DCL | Data from camera to lens |
| SCK (pin 13) | LCLK | Data transfer clock |
| **External power supply** | | |
| VCC | VBAT | AFD-USM, EMD drive power supply |
| GND | P-GND | V-BAT ground |

Figure 5: Pinout diagram.

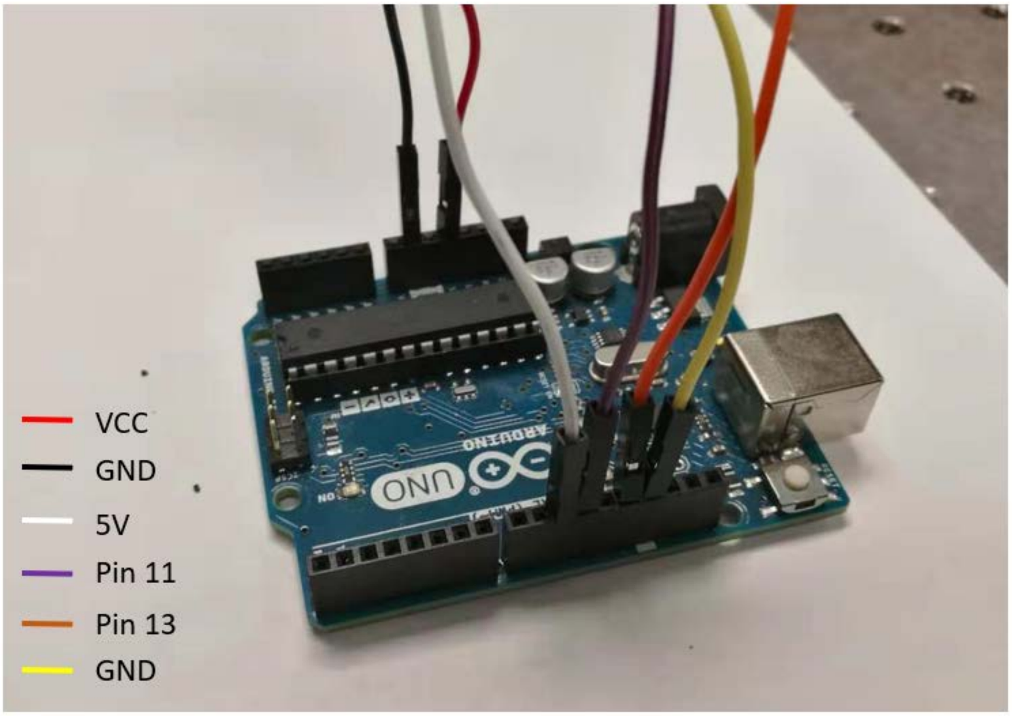


Figure 6: Wire connection with Arduino board.

**Step 4: Arduino Code for the photographic lens control**

- At first, we define mode and state of pins.

```
const int LogicVDD_Pin=10;
const int Cam2Lens_Pin=11;
const int Clock_Pin=13;
digitalWrite (Clock_Pin, LOW);
pinMode (LogicVDD_Pin, OUTPUT);
digitalWrite (LogicVDD_Pin, HIGH);
pinMode (Cam2Lens_Pin, OUTPUT);
digitalWrite (Cam2Lens_Pin, HIGH);
```

- Second, we initial the SPI (Serial Peripheral Interface) interface.

```
SPI.begin ( );
SPI.setBitOrder (MSBFIRST);
SPI.setClockDivider (SPI_CLOCK_DIV128);
SPI.setDataMode (SPI_MODE3);
```

- Third, we give an initial position of lens ring is 5000.

```
apAddr = 0;
```

```
focuserPosition = 5000;
IsMoving = false;
IsFirstConnect = ture;
```

- Next, we set the baud rate to 2000000.

```
Serial.begin (2000000);
```

- Finally, we initialize the lens.

```
void InitLens ( )
{
 SPI.transfer (0x??);
 delay (20) ;
 SPI.transfer (0x00);
 delay (20) ;
 SPI.transfer (0x??);
 delay (20) ;
 SPI.transfer (0x00);
 delay (20) ;
}
void loop ()
{
        if (Serial.available ( ) > 0)
        {
                Serial.readBytes (inStr, 5) ;
                targetStr = inStr ;
                targetPos = (targetStr).toInt ( ) ;
                offser = targetPos – focuserPosition ;
                x = highByte (offset) ;
                y = lowByte (offset) ;
                IsMoving = true ;
                SPI.transfer (0x??) ;
                delay (10) ;
                SPI.transfer (x) ;
                delay (10) ;
                SPI.transfer (y);
                delay (10) ;
```

```
        SPI.transfer (0) ;
    }
}
```

Note: For the commands of ??, one can

**Reference**

[1] http://web.media.mit.edu/~bandy/invariant/move_lens.pdf
