## Supplementary material 3 for "High-throughput functional characterization of visceral afferents by optical recordings from thoracolumbar and lumbosacral dorsal root ganglions"

### Instruction manual for GCaMP6f transient signal extraction from image sequences

#### 1 Software installation

We use MATLAB for data processing in this protocol. MATLAB is a programming platform designed specifically for engineers and scientists. It can directly express matrix and array mathematics[1].

MATLAB and the Image Processing Toolbox can be downloaded and installed following these two links: https://www.mathworks.com/ downloads/,
https://www.mathworks.com/products/image.html.

#### 2 Image alignment using the mutual information

To address the motion artifacts in our recorded image stacks, we first perform image alignment by maximizing the mutual information (MI) of different images. The positional shifts or rotations in-between different frames can be corrected in this process. We also note that this operation is not required if user's data set doesn't suffer from motion artifacts.

The operations of the reported MATLAB GUI for image alignment:

1) Open the file 'Image_alignment.m' in MATLAB. Click the green 'Run Section' icon. The GUI will appear like below.

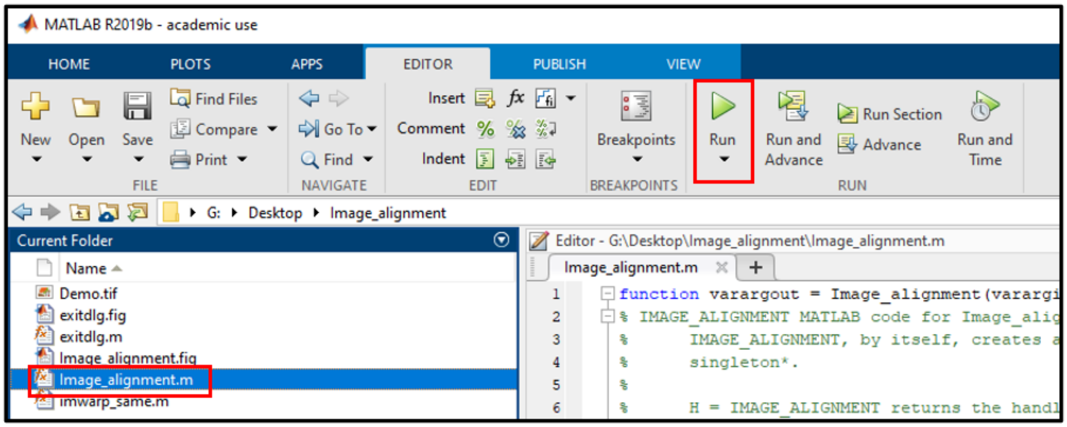

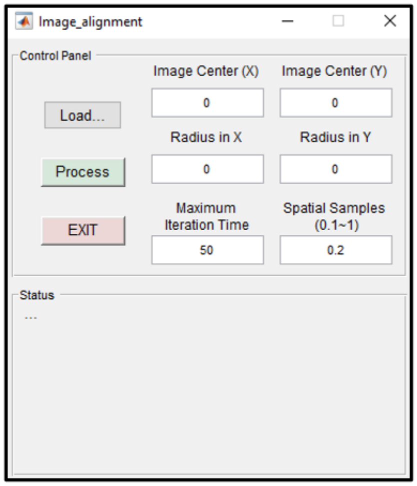


2) After opening the GUI, the first step is to load the original image stack into the MATLAB workspace. Here, we use the 'Demo_stack.tif' file as an example. First, click the 'Load…' button, a pop-up window will appear. Select the 'Demo_stack.tif' and click the 'Open' button.

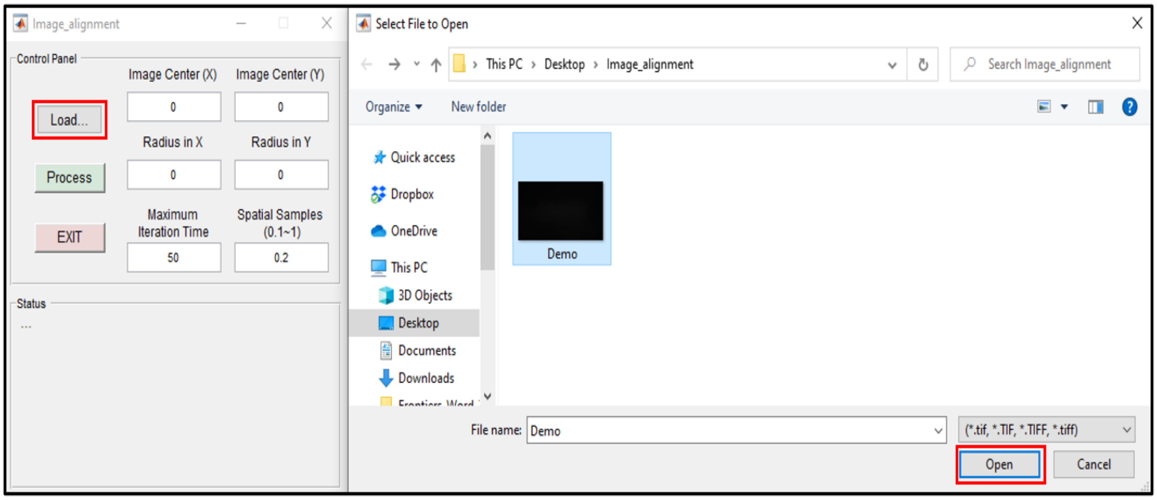


It will load the stack into the MATLAB workspace. When Step 1 is completed, the first frame of the stack will be displayed and the 'Status' panel will appear as follows:

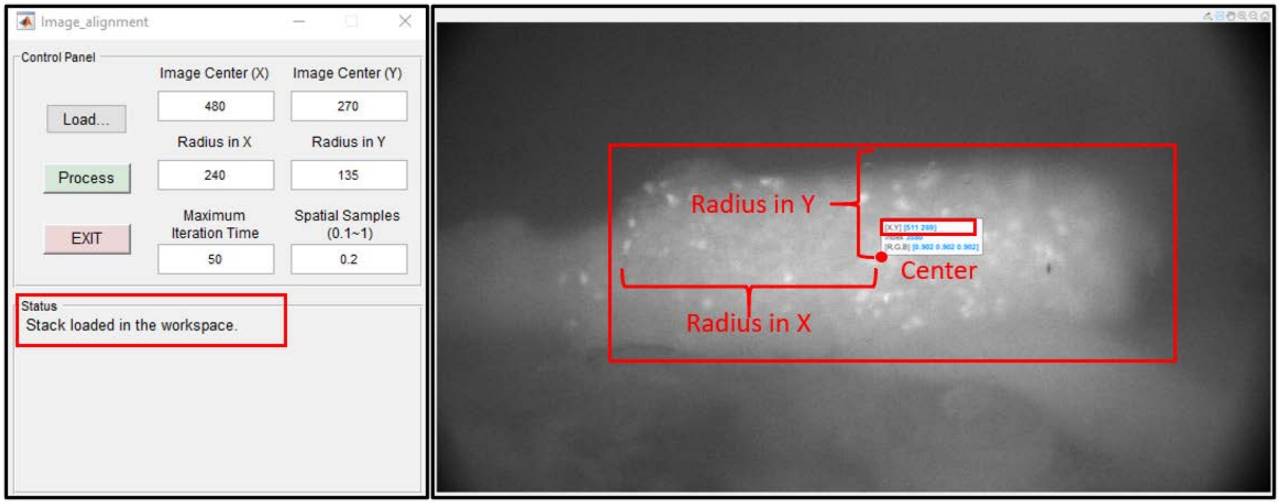

3) The second step is to perform image alignment. We use the parameters ‘Image Center (in X, Y)’ and ‘Radius (in X, Y)’ to represent the estimated the region of entire DRG (in pixels). In the alignment process, we use gradient decent optimizer to maximize the Mutual Information (MI). The parameter ‘Maximum Iteration Time’ defines the maximum number of iterations the optimizer performs at given settings. Similarly, the parameter ‘Spatial Samples’ represents the number of pixels that will be used for MI computation (1 means all pixels, 0.1 means 10% pixels). These parameters can be adjusted based on the magnification factor of the microscope setup and the specimens under testing. After clicking the ‘Process’ button, the alignment will be performed and the ‘Status’ panel will change to ‘Processing…’ as shown below.

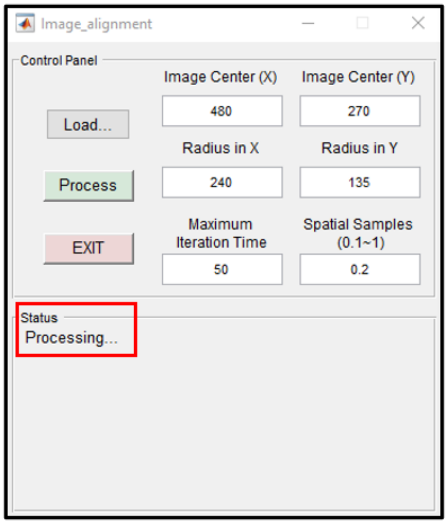


When the process is completed, an aligned frame will be displayed and the corrected stack will be saved under the same folder. Supplementary Video 1 compares the image difference before and after the alignment process .

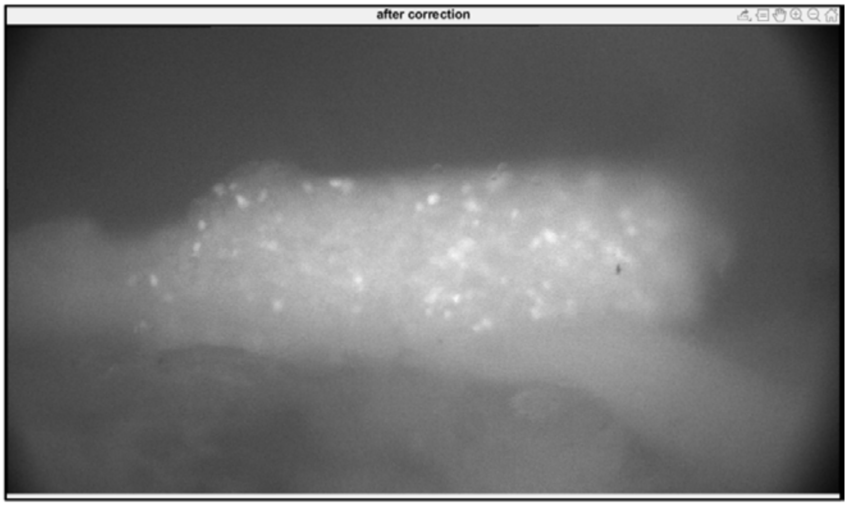


#### 3 Extracting GCaMP6f transient signal

A MATLAB graphical user interface (GUI) has also been provided to further extract the GCaMP6f transient signal from the aligned image stack. The first step is to load the image stack based on the user’s selection. In the second step, the morphological operations are applied to the image stack.

Regions with DRG signals can be identified by marker-based watershed segmentation. The third step is to extract all signals in the form of pixel intensity. The last step is to display and save all response curves.

The operations of this MATLAB GUI:

1) Open the file ‘neuron_detector.m’ in MATLAB. Click the green ‘Run Section’ icon. The GUI will appear like below.

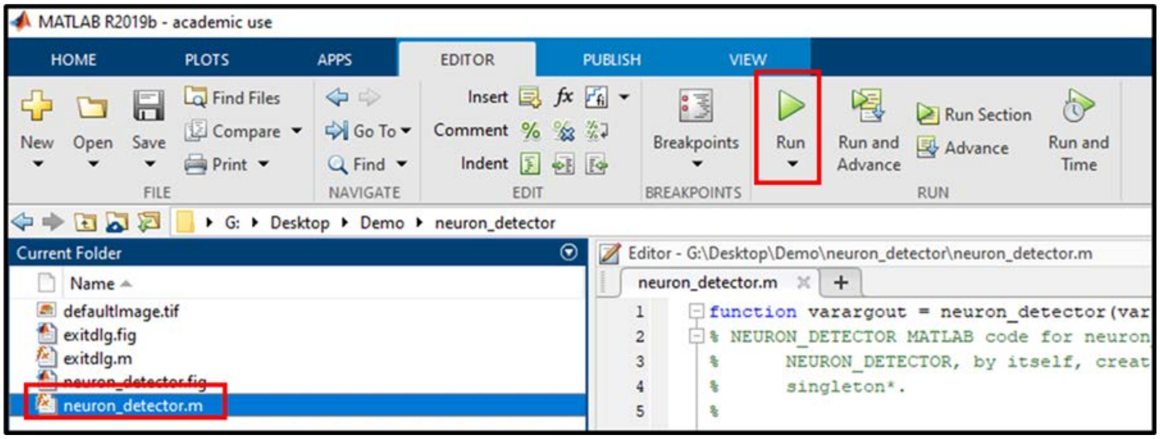


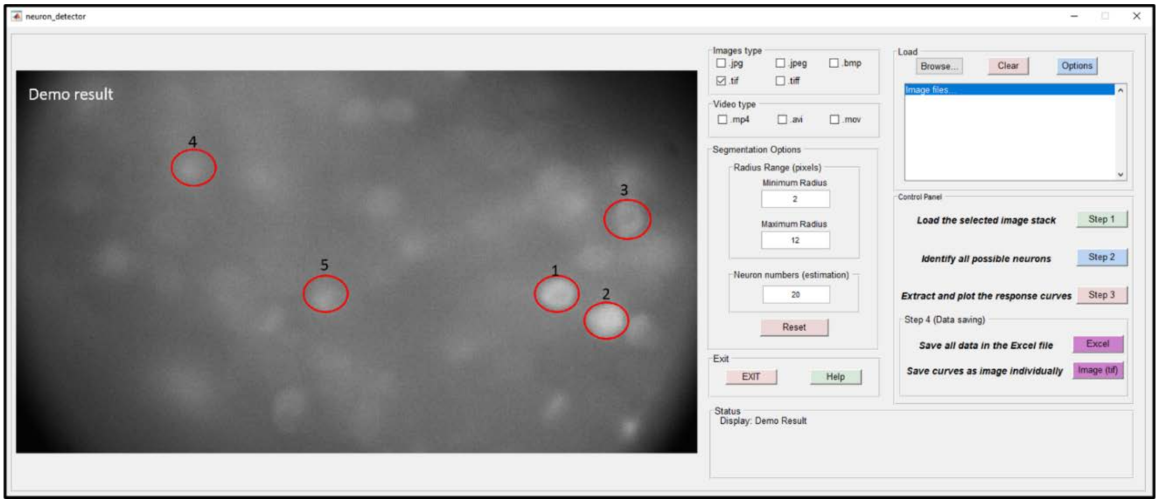


2) After opening the GUI, the first step of operations is to load the correct image stack into the MATLAB workspace. Our GUI is able to recognize multiple image formats and video formats. Here, we use the ‘Demo_stack.tif’ file as an example. First, select ‘.tif’ checkbox. Second, click the ‘Browse…’ button, and a pop-up window will appear. Select the ‘Demo_stack.tif’ and click the ‘Open’ button.

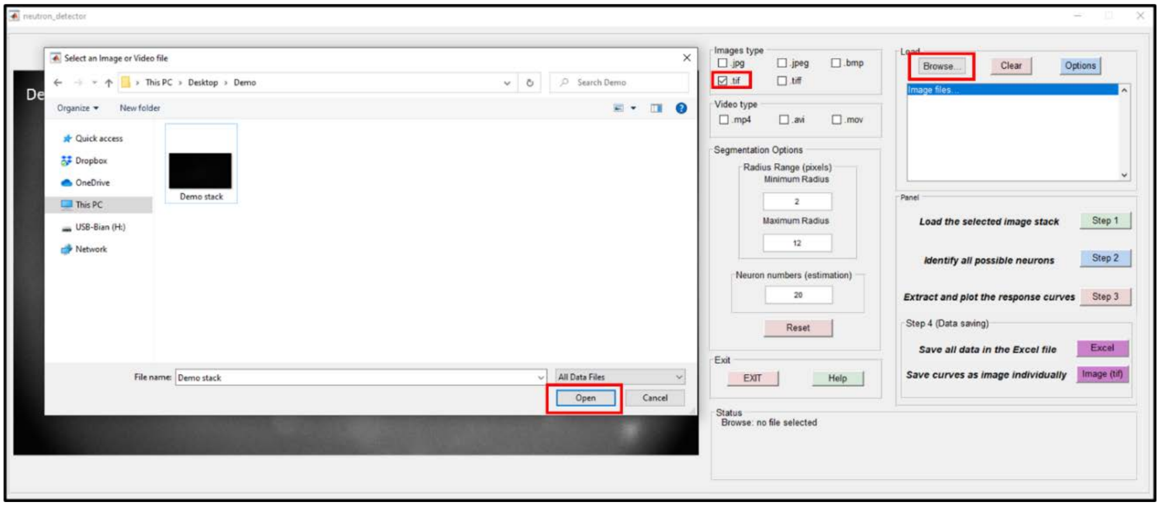


All tif files under the same file path will be listed in the GUI. In this example, the 'Demo_stack.tif' is the only tif file under the selected path. Select and left click the file's name. The first frame of the stack will appear in the GUI and the 'Status' panel will also list the file's name.

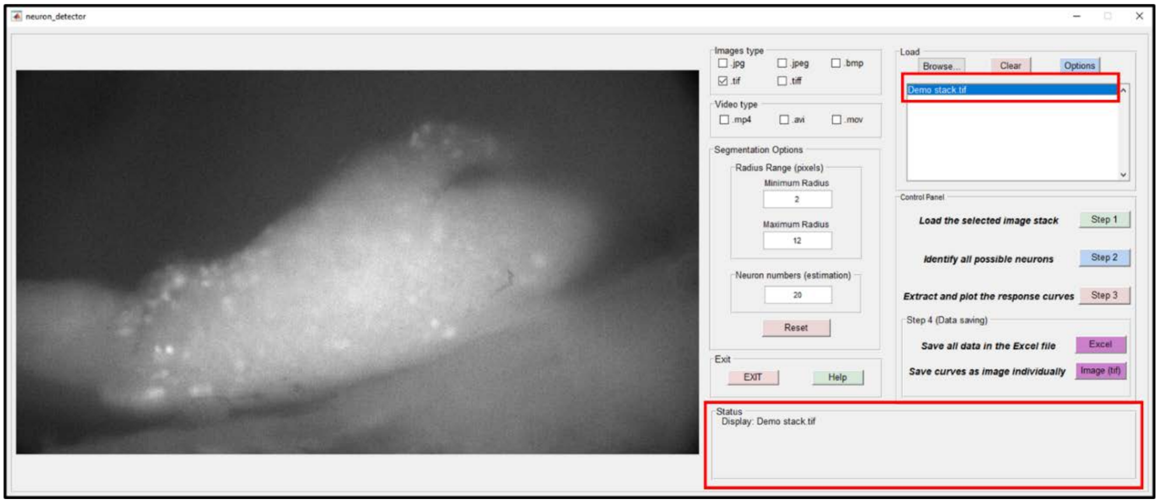


After selecting the tif file, click the 'Step 1' button. It will load the stack into the MATLAB workspace. The process bar will appear and the 'Status' panel will change to 'Loading…' as shown below.

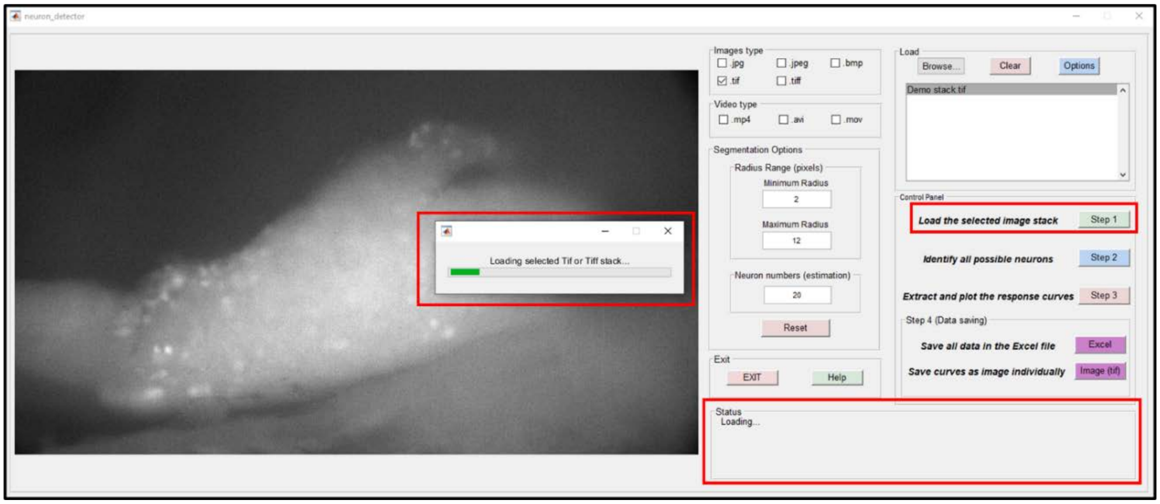


After the ‘Loading’ process, a ‘normalization’ operation will be applied to the image stack. Each frame will be divided by its mean intensity value to remove background fluctuation. Following the normalization operation, a variation map will be generated to show the intensity variations of the image stack.

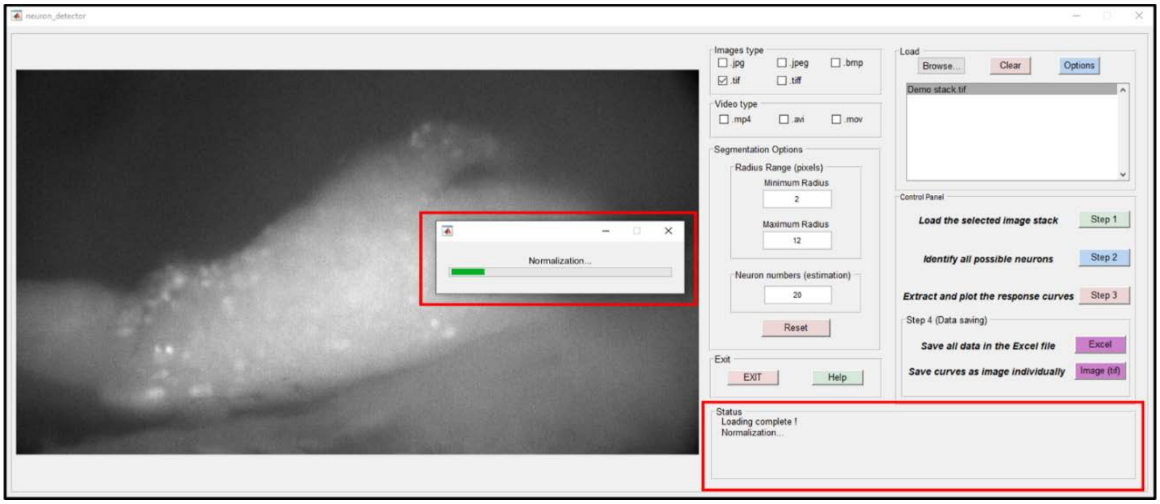


When Step 1 is completed, the ‘Status’ panel will appear as follows:

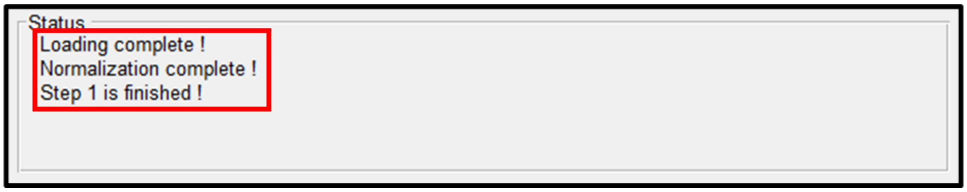


3) The second step of the operation is to identify all possible active neurons via the variation map. We use the parameters ‘Minimum Radius’ and ‘Maximum Radius’ to represent the estimated minimal and maximal radius of DRG neurons (in pixels). The identified regions outside the radius region will be excluded from the final result. The parameter ‘Neuron numbers’ represents the estimated number of positive neurons in the field of view. These three parameters can be adjusted based on the magnification

factor of the microscope setup and specimens under testing. After clicking the 'Step 2' button, the process bar will appear and the 'Status' panel will change to 'Start measuring…' as shown below.

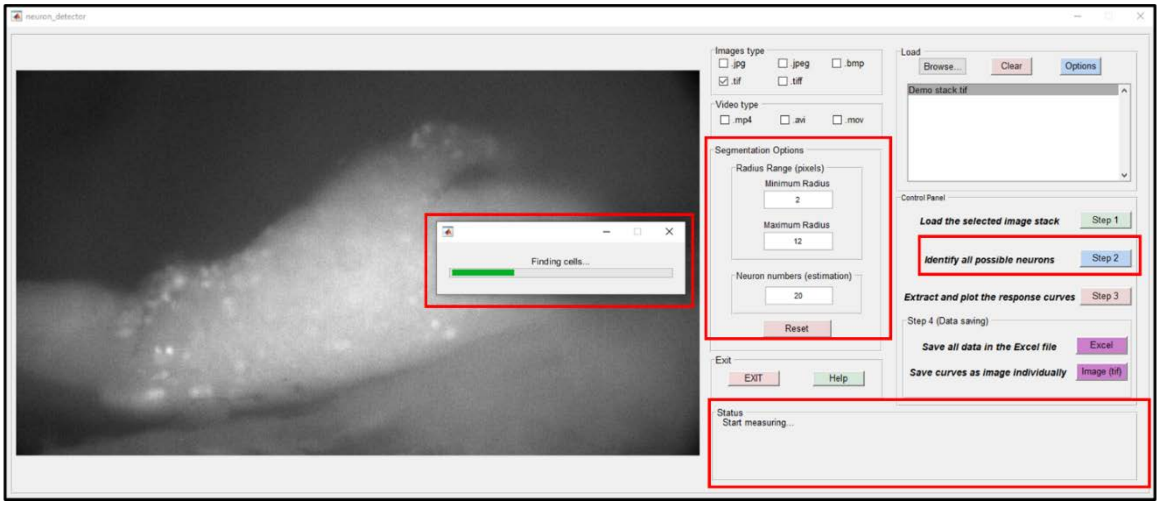


The 'Reset' button can adjust three parameters back to our default settings.

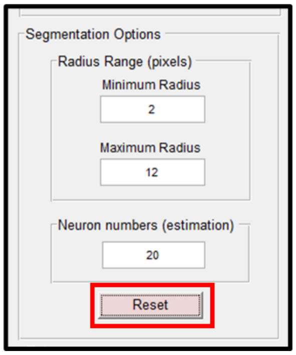


When Step 2 is completed, the image with labeled neurons will be shown in the GUI. The 'Status' panel will also list the number of possible neurons, and states 'Step 2 is finished!'.

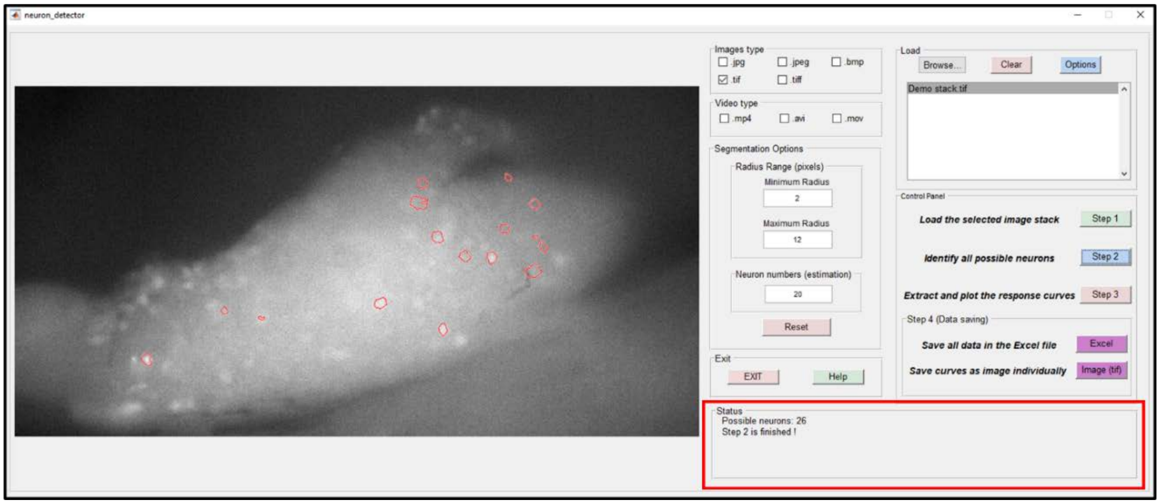

4) Step 3 of the operation is to extract and plot the response curves from the identified neurons. The average normalized pixel intensity of the labeled neuron is calculated to represent its response over time. After clicking the ‘Step 3’ button, the process bar and the ‘Status’ panel will appear as shown below.

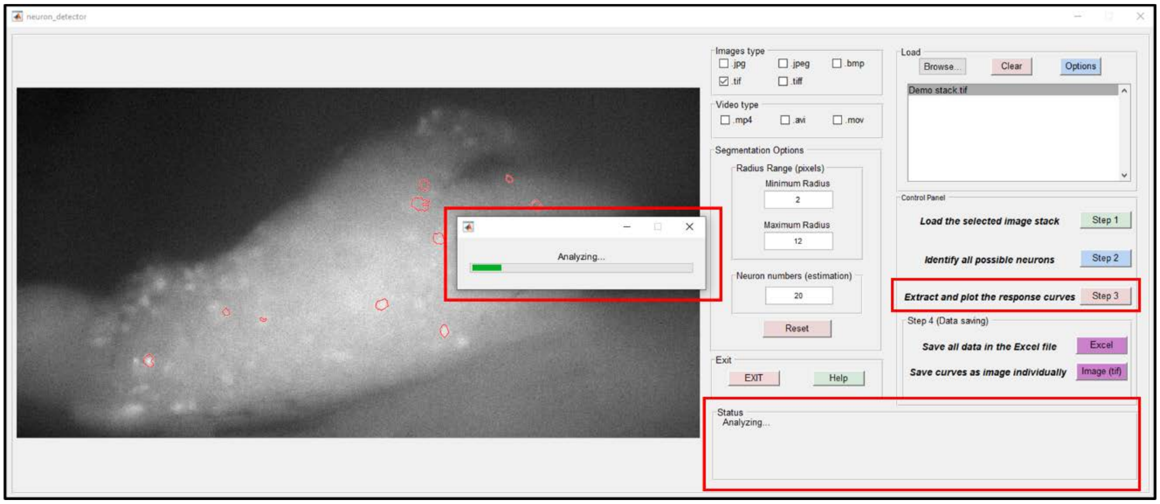


When Step 3 is completed, a full-screen figure with all response curves will appear as the figure shown below. The curves are ranked by their amplitude of variation. Some negative neurons are detected in this process. A better ranking strategy will be included in a later version of the protocol for removing negative responses.

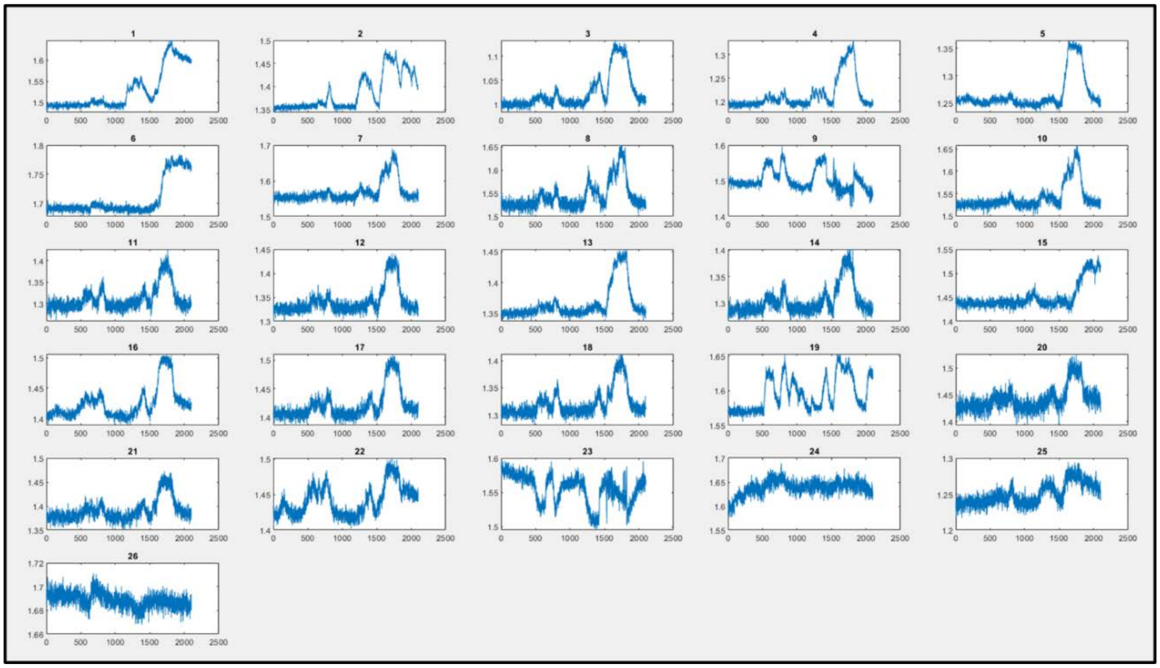

The ‘Status’ panel will change to ‘Step 3 is finished!’, as shown below.

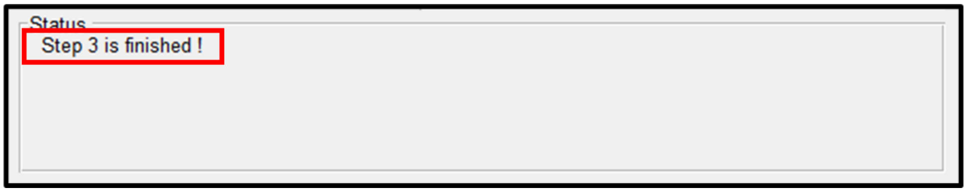


5) Step 4 of the operation is to save the response curve data. There are two options shown below. For the first option, user can save all data in an Excel file: click the 'Excel' button -> select the desired saving folder and click the 'Select Folder' button.

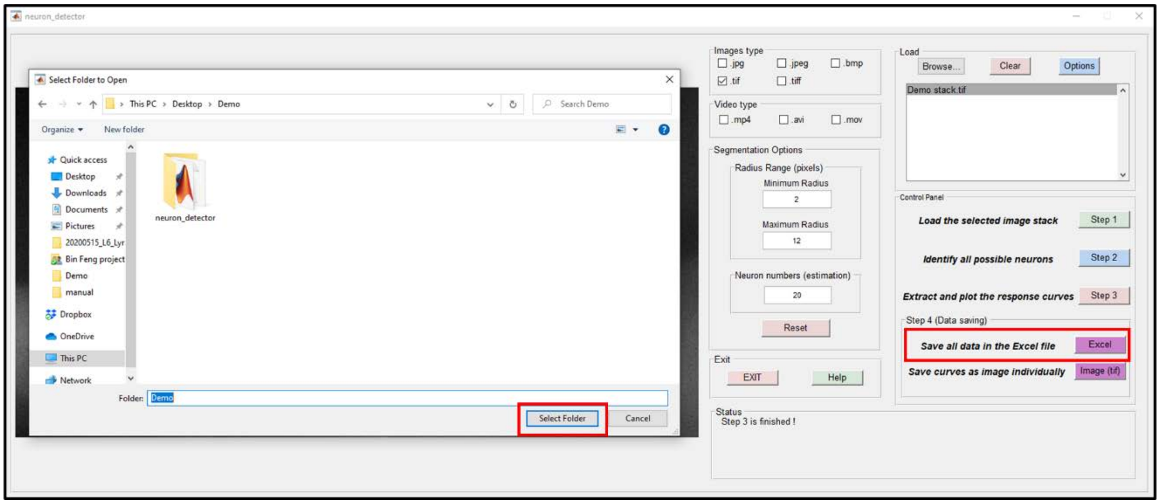


When the saving process is completed, the 'Status' panel will list the file path as shown below. A figure with all response curves (same as the Step 3 result) will also be saved in the same folder.

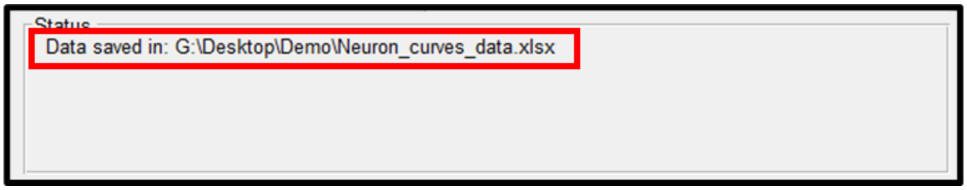


In this example, the 'Neuron_curves_data .xlsx' Excel file will contain all neurons' response curves:

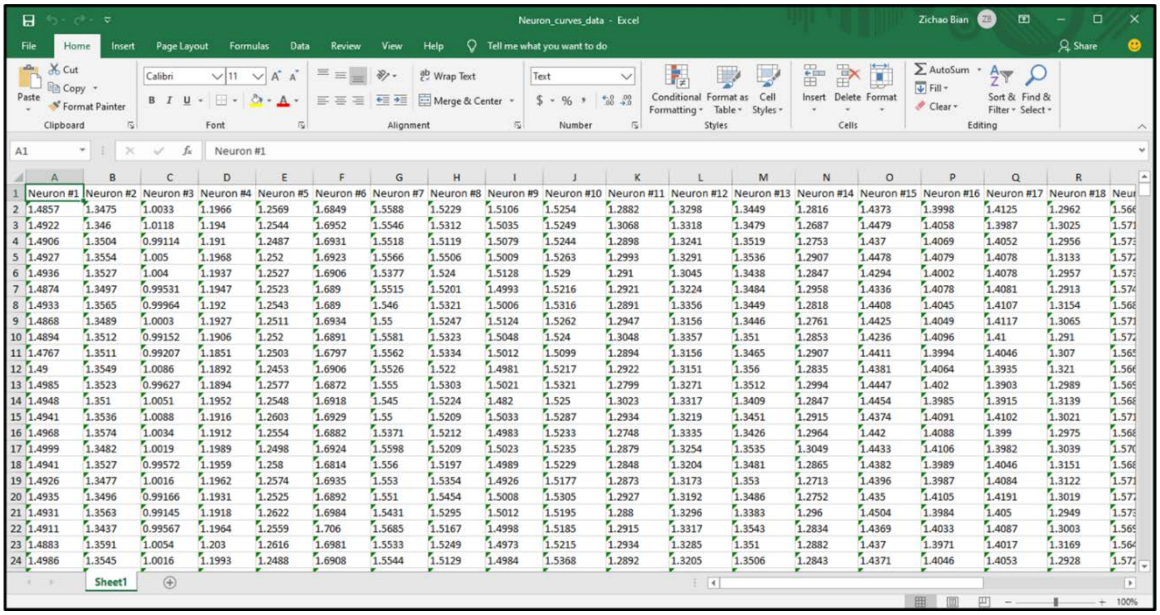


For the second option, user can save every curve with its corresponding neuron's location as a tif image file. All the tif files will be merged into one tif stack. After clicking the 'Image (tif)' button, the dialog window will appear as the figure below.

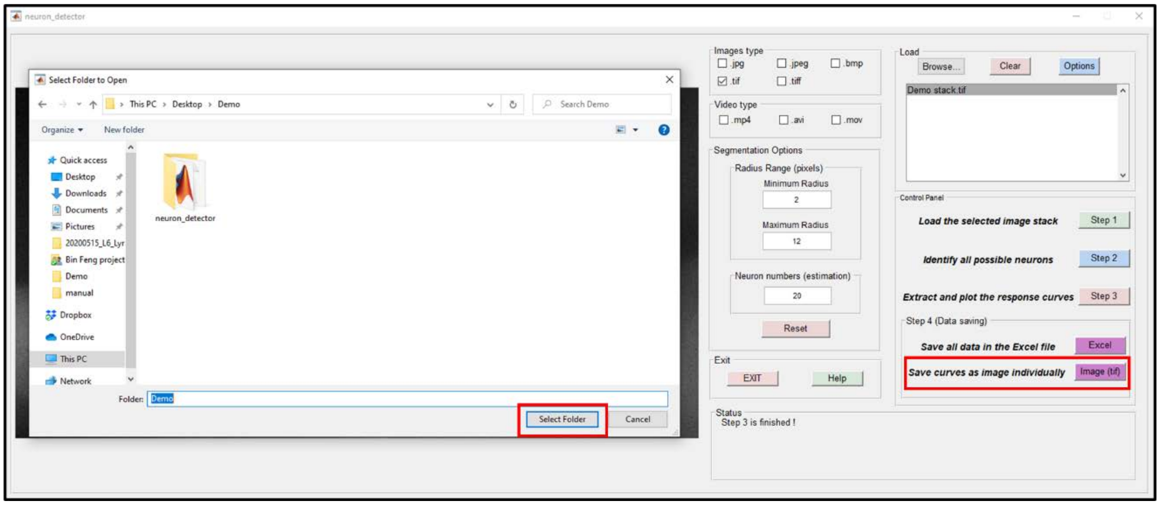


When the saving process is completed, the 'Status' panel will list the file path as shown below.

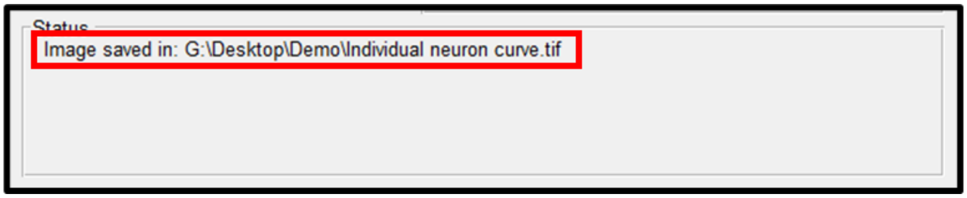


The 'Individual neuron curve.tif' stack contains all neurons' curves and the corresponding locations. The frame number of this stack equals the possible neurons' number as shown below.

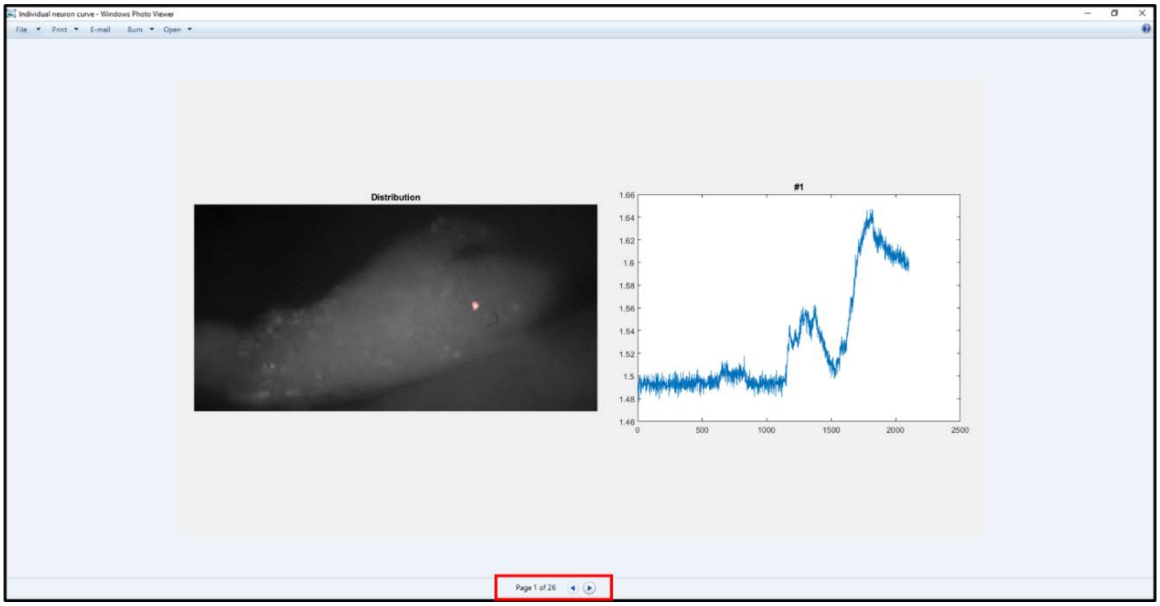


6) When all operations are completed, user can click the ‘EXIT’ button and select ‘Yes’ to exit the MATLAB GUI.

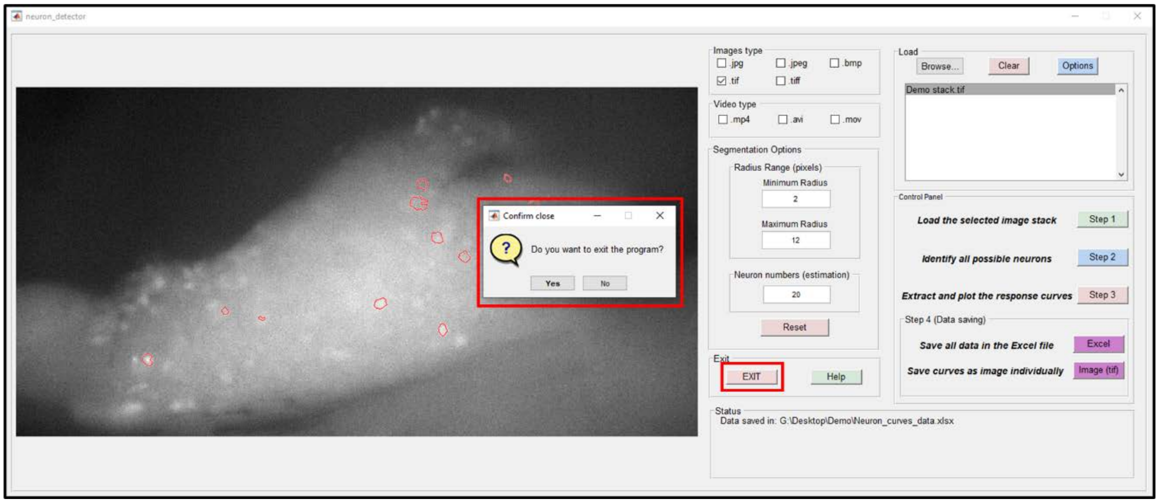


#### 4 Reference:

1. “What Is MATLAB?” MATLAB & Simulink, 2020, www.mathworks.com/discovery/what-is-matlab.html.
