## Supplementary material 4 for "High-throughput functional characterization of visceral afferents by optical recordings from thoracolumbar and lumbosacral dorsal root ganglions"

### Control of the colorectal distension device

**Step 1: Wire Connection**

The valve system for graded colorectal distension and luminal shear stimuli is control by an Arduino Uno board (ATmega328P) that receives commands from a computer. In this system, we use the N-Channel Metal-Oxide-Semiconductor Field-Effect Transistor (MOSFET) as switch to control the on-off state of the valve. The drain of the MOSFET is connected with one end of solenoid valve, the other end is connected to a 12 V DC power supply. The source of the MOSFET is connected to the GND of the Arduino board. The gate of the MOSFET is concatenated with the digital pin of the Arduino board, the logical level of which is determined by the computer command. By adjusting the logical value (HIGH/LOW) of the digital pins, the corresponding valve will be set as on or off. The current system is capable of controlling 7 solenoid valves sufficient to fulfill the demand of experiment. The schematic connection of valve 1 is shown as an example in the figure below.

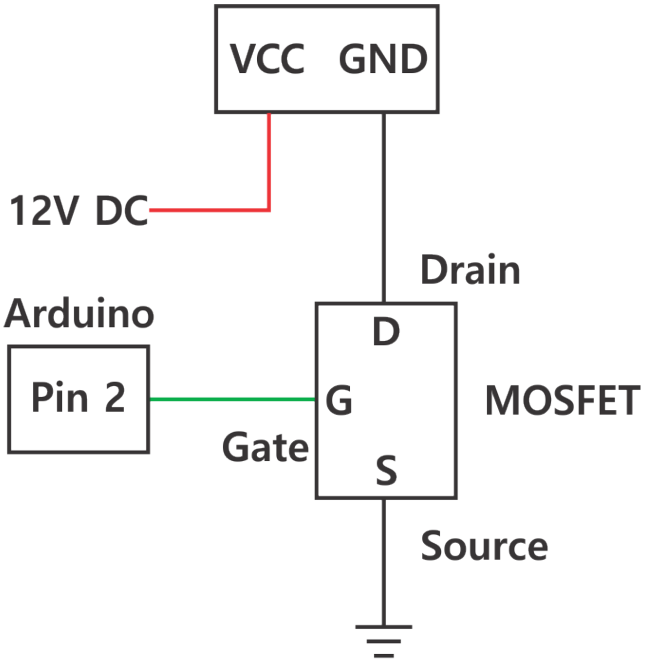


Figure 1: Valve 1 connection diagram.

**Step 2: Arduino Code for the valves control**

- First, we define the mode and state of different pins.

```
int valve1=2;
int valve2=3;
int valve3=4;
int valve4=5;
```

```
int valve5=6;
int valve6=7;
int valve7=8;
pinMode (valve1, OUTPUT);
pinMode (valve2, OUTPUT);
pinMode (valve3, OUTPUT);
pinMode (valve4, OUTPUT);
pinMode (valve5, OUTPUT);
pinMode (valve6, OUTPUT);
pinMode (valve7, OUTPUT);
```

Note: Avoid using digital pin 0 and pin 1 as the voltage control for the valves, they are used for the serial communication.

- Next, we set the baud rate to be 2000000.

```
Serial.begin (2000000);
```

- Finally, we define the ‘switch’ condition for each valve.

```
void loop ()
{
        if (Serial.available ( ) > 0)
        {
                char cx = Serial.read();
                    switch (cx) {
                    case 'a':
                    // Turn on the valve 1
                    digitalWrite(valve1, HIGH);
                    digitalWrite(valve2, LOW);
                    digitalWrite(valve3, LOW);
                    digitalWrite(valve4, LOW);
                    digitalWrite(valve5, LOW);
                    digitalWrite(valve6, LOW);
                    digitalWrite(valve7, LOW);
                    break;

                    case 'b':
```

```
// Turn on the valve 2
 digitalWrite(valve1, LOW);
 digitalWrite(valve2, HIGH);
 digitalWrite(valve3, LOW);
 digitalWrite(valve4, LOW);
 digitalWrite(valve5, LOW);
 digitalWrite(valve6, LOW);
 digitalWrite(valve7, LOW);
 break;

 case 'c':
 // Turn on the valve 3
 digitalWrite(valve1, LOW);
 digitalWrite(valve2, LOW);
 digitalWrite(valve3, HIGH);
 digitalWrite(valve4, LOW);
 digitalWrite(valve5, LOW);
 digitalWrite(valve6, LOW);
 digitalWrite(valve7, LOW);
 break;

 case 'd':
 // Turn on the valve 4
 digitalWrite(valve1, LOW);
 digitalWrite(valve2, LOW);
 digitalWrite(valve3, LOW);
 digitalWrite(valve4, HIGH);
 digitalWrite(valve5, LOW);
 digitalWrite(valve6, LOW);
 digitalWrite(valve7, LOW);
 break;

 case 'e':
 // Turn on the valve 5
```

```
digitalWrite(valve1, LOW);
digitalWrite(valve2, LOW);
digitalWrite(valve3, LOW);
digitalWrite(valve4, LOW);
digitalWrite(valve5, HIGH);
digitalWrite(valve6, LOW);
digitalWrite(valve7, LOW);
break;

case 'f':
// Turn on the valve 6
digitalWrite(valve1, LOW);
digitalWrite(valve2, LOW);
digitalWrite(valve3, LOW);
digitalWrite(valve4, LOW);
digitalWrite(valve5, LOW);
digitalWrite(valve6, HIGH);
digitalWrite(valve7, LOW);
break;

case 'g':
// Turn on the valve 7
digitalWrite(valve1, LOW);
digitalWrite(valve2, LOW);
digitalWrite(valve3, LOW);
digitalWrite(valve4, LOW);
digitalWrite(valve5, LOW);
digitalWrite(valve6, LOW);
digitalWrite(valve7, HIGH);
break;

case 'A':
// Turn off all valves
```

```
            digitalWrite(valve1, LOW);
            digitalWrite(valve2, LOW);
            digitalWrite(valve3, LOW);
            digitalWrite(valve4, LOW);
            digitalWrite(valve5, LOW);
            digitalWrite(valve6, LOW);
            digitalWrite(valve7, LOW);
            break;
    }
}
```
